# N-terminal acetylation controls multiple functional aspects of the influenza A virus ribonuclease PA-X

**DOI:** 10.1101/2023.12.01.569683

**Authors:** Raecliffe E. Daly, Cynthia Y. Feng, Charles Hesser, Idalia Myasnikov, Marta Maria Gaglia

## Abstract

To counteract host antiviral responses, influenza A virus triggers a global reduction of cellular gene expression, a process termed “host shutoff.” A key effector of influenza A virus host shutoff is the viral endoribonuclease PA-X, which degrades host mRNAs. While many of the molecular determinants of PA-X activity remain unknown, a previous study found that N-terminal acetylation of PA-X is required for its host shutoff activity. However, it remains unclear how this co-translational modification promotes PA-X activity. Here, we report that PA-X N-terminal acetylation has two functions that can be separated based on whether acetylation occurs on the first amino acid, the initiator methionine, or the second amino acid following initiator methionine excision. Modification at either site is sufficient to ensure PA-X localization to the nucleus. However, modification of the second amino acid is not sufficient for host shutoff activity of ectopically expressed PA-X, which specifically requires N-terminal acetylation of the initiator methionine. Interestingly, during infection N-terminal acetylation of PA-X at any position results in host shutoff activity, suggesting that additional factors during infection can augment the host shutoff activity of PA-X. Our studies thus uncover a multifaceted role for PA-X N-terminal acetylation in regulation of this important immunomodulatory factor.

**Importance:** Influenza A viruses pose a significant threat to human health in the form of seasonal epidemics and recurrent pandemics, leading to large burdens on the healthcare system. During infection, host immune and inflammatory responses have a key role in disease outcome. They are needed to clear the virus, but when excessive they can cause lung damage. Influenza A viruses encode several factors that modulate the host immune response, influencing viral replication and pathogenesis. In this study, we focused on one of these influenza factors, PA-X, which destroys cellular mRNAs and reduces cellular gene expression (a phenomenon called “host shutoff”) to control immune responses. This protein is modified with an acetylation at its N-terminus, and this modification is needed for its activity, but it has remained unclear why. Our results demonstrate that N-terminal acetylation of PA-X is needed for multiple aspects of its function. It ensures that PA-X goes to the nucleus, where it accesses its RNA targets, but it also separately contributes to host shutoff activity. However, for host shutoff activity, the specific location of the modification matters, whereas for entry in the nucleus it does not. For full activity, the modification must be placed on the very first amino acid of the protein, and this amino acid must be methionine. These findings uncover how influenza A viruses exploit a widespread protein modification to support the host shutoff activity of one of its important immunomodulatory proteins.

## Introduction

Although influenza A viruses have a relatively small genome of approximately 13.5 kb, multiple influenza proteins are involved in modulating the host immune response. Of the influenza immunomodulatory proteins identified to date, PA-X has emerged as a key factor in regulating gene expression and innate immune responses. Studies using viruses engineered to lack PA-X have revealed that by limiting innate immune and inflammatory responses during infection, PA-X reduces inflammation-induced pathology(1, 2). Immune regulation by PA-X is the result of its crucial role in influenza host shutoff, i.e. the global reduction of gene expression during infection(1, 3, 4). However, since PA-X was only discovered in 2012(1), studies of this protein are still limited.

Previous studies have focused on how PA-X drives host shutoff and have shown that PA-X has endoribonuclease (endoRNase) activity(1, 4, 5). PA-X fragments host mRNAs, while largely sparing viral RNAs(4, 6). PA-X discriminates host and viral mRNAs in part based on its sequence preference(6), as it preferentially cleaves GCUG tetramers within hairpin loops, which are enriched in the human but not the influenza transcriptome(6). PA-X also preferentially degrades mRNAs that are spliced, whereas most viral mRNAs are not(5). While these previous studies have dissected the specificity of RNA targeting by PA-X, we still have limited information on how PA-X functions. We and others have shown that it accumulates in the nucleus, and that nuclear localization is required for activity(4, 7). We also previously reported that PA-X is rapidly turned over, with protein half-lives ranging from ∼30 minutes to ∼3.5 hours depending on the strain(8). Whether and how PA-X activity is modulated in other ways in cells remains unclear.

To date, only one protein modification of PA-X has been reported. Oishi et al. found that PA-X is N-terminally acetylated by the host N-terminal acetyltransferase complex B (NatB)(9). N-terminal acetylation is a highly abundant co-translational protein modification in cells, present on 80-90% of the human proteome(10–13). It is catalyzed by several Nat complexes (NatA-NatF), which differ in their subunit composition and substrate specificity(14, 15). NatA, NatB, and NatC are responsible for the majority of N-terminal acetylation events in eukaryotes(15). Their substrate specificity is mainly determined by the identity of the second N-terminal residue of the target protein(14) (Fig. 1A). NatB and NatC acetylate nascent polypeptides on the very first translated amino acid, the initiator methionine, whereas NatA modifications occur following the excision of the initiator methionine by methionine aminopeptidases. PA-X starts with an ME-sequence that is highly conserved among influenza isolates(9). Therefore, almost all natural variants are modified by NatB(9). Despite the fact that N-terminal acetylation is a common modification, only a handful of studies have investigated the effect of N-terminal acetylation on protein function(15, 16). Nonetheless, N-terminal acetylation has been implicated in multiple processes, including the control of protein stability(17–21), protein folding(22–24), protein-protein interactions(25–30), and subcellular targeting(31–34). While Oishi et al. showed that PA-X requires this modification to carry out host shutoff(9), it remains unknown how N-terminal acetylation supports PA-X host shutoff activity.

**Fig 1.**
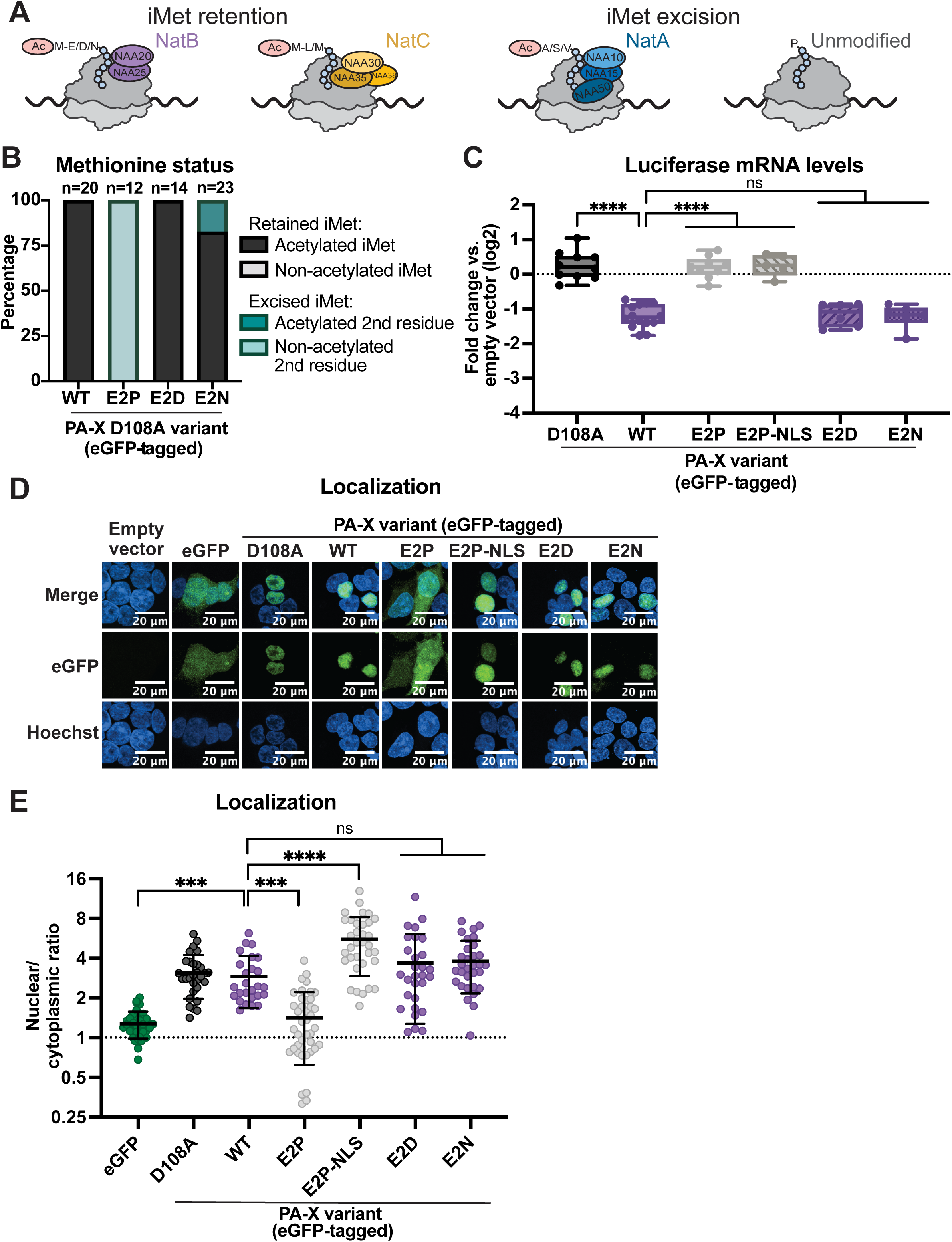
N-terminal acetylation of PA-X is required for nuclear localization. A) Diagram showing NatA-C complexes and the proteins they acetylate depending on the first two amino acids, and whether they require excision of the initiator methionine (iMet). Based on the described properties of NatA-C, WT PR8 PA-X and the PR8 PA-X E2 mutants E2D and E2N should be modified by NatB (purple), PA-X E2L and E2M mutants should be modified by NatC (yellow), and PA-X E2A, E2S, and E2V mutants should be modified by NatA (blue). The PA-X E2P mutant should be unmodified. B-E) HEK293T cells were transfected for 24 hours with empty vector, WT PR8 PA-X-eGFP, the catalytically inactive PA-X D108A-eGFP mutant, the indicated PR8 PA-X-eGFP mutants carrying changes in the second amino acid in the WT or D108A PR8-PA-X-eGFP background, or an eGFP-tagged PR8 PA-X E2P mutant carrying an SV40 nuclear localization sequence (PA-X E2P-NLS-eGFP). B) eGFP-tagged proteins were isolated by immunoprecipitation and peptides were analyzed by nanoLC-MS/MS for iMet retention and N-terminal acetylation. N above stacked bars indicate total detected peptides for each variant over two experiments. C) mRNA levels of a co-transfected luciferase reporter were measured by RT-qPCR and normalized to 18S rRNA. Levels are plotted as fold change relative to vector transfected cells in log2 scale. D) Confocal microscopy was used to image empty vector, eGFP, PA-X D108A-eGFP (D108A), PA-X-eGFP (WT), PA-X E2P-eGFP (E2P), PA-X E2P-NLS-eGFP (E2P-NLS), PA-X E2D-eGFP (E2D) and PA-X E2N-eGFP (E2N). Nuclei were stained with Hoechst. E) Nuclear/cytoplasmic ratios were calculated by measuring the mean fluorescence intensity of the nuclear and cytoplasmic compartments using ImageJ. Hoechst staining was used as a guide to generate regions of interest for the nucleus. Autofluorescence was used as a guide to generate cytoplasmic regions of interest. Nuclear/cytoplasmic ratios represent the following localization: <1: protein is primarily cytoplasmic, ∼1: protein is diffuse throughout the cell, >1: protein is primarily nuclear. For all experiments, some conditions were carried out in parallel to experiments in Figures 2 and 3. Therefore, control conditions (Empty vector, PA-X D108A, WT PA-X, and eGFP) are identical. N≥3 for all experiments except the MS analysis in panel A. For microscopy, representative images are shown. In nuclear/cytoplasmic ratio quantitation, each point represents 1-4 eGFP-positive cells, with 10-15 eGFP-positive cells quantified per condition in each biological replicate. ns = p > 0.05, *** = p < 0.001, **** = p < 0.0001, ANOVA with Dunnett’s multiple comparison test.

A complication in studying PA-X N-terminal acetylation is that PA-X shares its N terminus with the polymerase acidic (PA) subunit of the influenza RNA polymerase, a protein that is essential for viral replication(1, 35). This is because PA-X is generated from the same mRNA as PA (the mRNA transcribed from the influenza genomic segment 3) through ribosomal frameshifting after translation of the 191^st^ amino acid of PA(1, 35). Therefore, mutations or acetylase knockdown/knockouts that alter PA-X N-terminal acetylation also impact PA N-terminal acetylation(9). Since PA also requires this modification for its function, again due to an unknown mechanism of regulation(9), the impact of PA-X N-terminal acetylation during infection was not separated from that of PA modification in the initial study from Oishi et al.(9).

Here, we report that N-terminal acetylation supports PA-X activity by two separate mechanisms. N-terminal acetylation is needed for PA-X nuclear localization(4, 7, 36). We found that PA-X localizes to the nucleus when modified by Nat complexes that add the modification to either the initiator methionine or the second residue following initiator methionine excision. However, in the absence of viral infection, the localization alone is not enough to confer host shutoff activity, and PA-X specifically requires modification of the initiator methionine to downregulate RNA levels in cells. This result suggests that N-terminal acetylation on the initiator methionine has a separate role in supporting PA-X host shutoff activity. To our surprise, we also observed that PA-X mutants modified at the second amino residue downregulate transcripts during infection, even though they are largely inactive when PA-X is ectopically expressed. This result indicates that PA-X activity is somehow potentiated during influenza A virus infection. The stimulation of PA-X activity during infection does not appear to be due to a single viral protein, but likely to another effect of influenza infection that remains to be determined.

## Results

### N-terminal acetylation is required for PA-X nuclear localization, but nuclear localization in the absence of acetylation is not sufficient for host shutoff activity

Previous studies using the influenza A/WSN/33 H1N1 (WSN) virus have shown that N-terminal acetylation of PA-X by the NatB complex is important for its host shutoff activity in mammalian cells(9). When the N terminus of WSN PA-X is mutated from ME-to MP-to produce non-acetylated PA-X proteins(15, 37) (Fig. 1A), the shutoff activity of WSN PA-X is lost(9). One of the established functions of this modification is control of subcellular localization(31–34) and we and others have reported that PA-X must localize to the nucleus to downregulate mRNAs(4, 7, 36). To determine whether non-acetylated mutants lose activity because of primarily cytoplasmic localization, we compared the subcellular localization of ectopically expressed wild-type (WT) PA-X from the influenza A/Puerto Rico/8/1934 H1N1 virus (PR8) and the non-acetylated PR8 PA-X E2P mutant, both tagged at the C-terminus with eGFP, using confocal microscopy. We used mass spectrometry to confirm that the WT protein is 100% acetylated, while the E2P mutant is not acetylated (Fig. 1B). Moreover, to confirm that the PR8 PA-X E2P mutant loses the ability to reduce gene expression like the WSN PA-X E2P mutant, we co-transfected cells with PA-X and a luciferase reporter and measured luciferase RNA levels by qPCR, as we have done in previous studies(4, 5). As expected, the PA-X E2P mutant did not downregulate luciferase mRNA levels, similarly to the catalytically inactive PA-X D108A mutant (Fig. 1C)(4, 5). We then imaged the localization of eGFP-tagged WT PA-X, PA-X D108A, and PA-X E2P, as well as eGFP alone (Fig. 1D). To quantify PA-X localization, we computed the ratio of the mean fluorescence of eGFP in the nucleus vs. the cytoplasm from the confocal microscopy images using Image J/Fiji(38, 39). Ratios lower than 1 indicate primarily cytoplasmic localization, ratios close to 1 indicate diffuse cellular localization, and values greater than 1 indicate primarily nuclear localization(38). eGFP alone showed an average nuclear/cytoplasmic ratio of 1.28, indicating diffuse localization throughout cells, whereas WT PA-X was predominantly nuclear, with an average nuclear/cytoplasmic ratio of 2.92, consistent with previous studies and with the images (Fig. 1D,E)(4, 7, 36). Interestingly, PA-X E2P was more diffusely localized than WT PA-X and had an average nuclear/cytoplasmic ratio of 1.42, similar to eGFP (Fig. 1D,E). These results suggest that N-terminal acetylation is required for the nuclear accumulation of PA-X, which could explain its loss of activity.

To ensure that the differences between WT PA-X and PA-X E2P were due to acetylation and not the sequence change, we also generated two mutants that have a different amino acid sequence but should still be modified by NatB like WT PA-X, PA-X E2D and PA-X E2N (Fig. 1A). In both the WSN(9) and the PR8 (Fig. 1C) background, these NatB modified mutants had similar host shutoff activity to WT PA-X, indicated by the ability to reduce expression of a coexpressed luciferase mRNA. Consistent with this result, PA-X E2D and PA-X E2N also had similar localization to WT PA-X, with nuclear/cytoplasmic ratios of 3.69 and 3.79, respectively (Fig. 1D,E). These results confirm that the differences seen between WT PA-X and PA-X E2P are due to changes in N-terminal acetylation.

To determine whether the decreased activity of PA-X E2P was solely due to its change in subcellular localization, we fused PA-X E2P to the canonical SV40 nuclear localization signal(40) to restore nuclear localization (Fig. 1D,E). Despite showing the expected nuclear localization (nuclear/cytoplasmic ratio = 5.56), PA-X E2P-NLS remained unable to downregulate luciferase mRNA levels (Fig. 1C). This result indicates that restoring the nuclear localization of an unmodified PA-X is not enough to restore its host shutoff activity. Therefore, we conclude that to downregulate RNA levels in cells, PA-X also requires N-terminal acetylation for a second function.

### N-terminal acetylation at any position is sufficient for PA-X localization to the nucleus, but not for host shutoff activity

To further explore the relationship between PA-X N-terminal acetylation, nuclear localization, and host shutoff activity, we mutated the second residue of PA-X to alter the Nat complex that modifies it (Fig. 1A). We chose to mutate PA-X rather than knock out the Nat complexes because 80-90% of the human proteome is N-terminally acetylated during translation, and knock-down of the complexes may have pleiotropic effects(10, 11, 13, 41). We first tested the E2A, E2S, and E2V mutations, which lead to N-terminal acetylation by NatA (Fig. 1A). Unlike NatB-modified proteins, which are N-terminally acetylated at the initiator methionine, NatA-modified proteins are N-terminally acetylated at the second residue following methionine excision(15, 42, 43) (Fig. 1A,). A previous study showed that, like the predominantly non-acetylated WSN PA-X E2P mutant, the NatA-modified WSN PA-X E2A mutant loses its host shutoff activity(9), suggesting the initiator methionine needs to be retained for PA-X function. Using mass spectrometry, we confirmed acetylation of these three mutants. PA-X E2A and E2S were 100% N-terminally acetylated at the second position following methionine excision, as expected (Fig. 2A). PA-X E2V showed a more varied pattern: the initiator methionine was only excised in about 80% of peptides (16 out of 21), and only about 90% (14 out of 16) of these were N-terminally acetylated. This is consistent with previous reports of partial N-terminal acetylation for proteins that start with MV-(13, 43). Like WSN PA-X E2A, PR8 PA-X E2A, E2S, and E2V had no apparent host shutoff activity in cells (Fig. 2B). Interestingly, despite losing the ability to downregulate mRNA levels, we saw that PA-X E2A, E2S, and E2V had a similar localization to WT PA-X, and primarily accumulated in the nucleus (Fig. 2C,D). Taken together, these data suggest that N-terminal acetylation at any position and at high levels is sufficient for PA-X nuclear localization, but that the differences between NatA and NatB modifications are important for host shutoff activity.

**Fig. 2.**
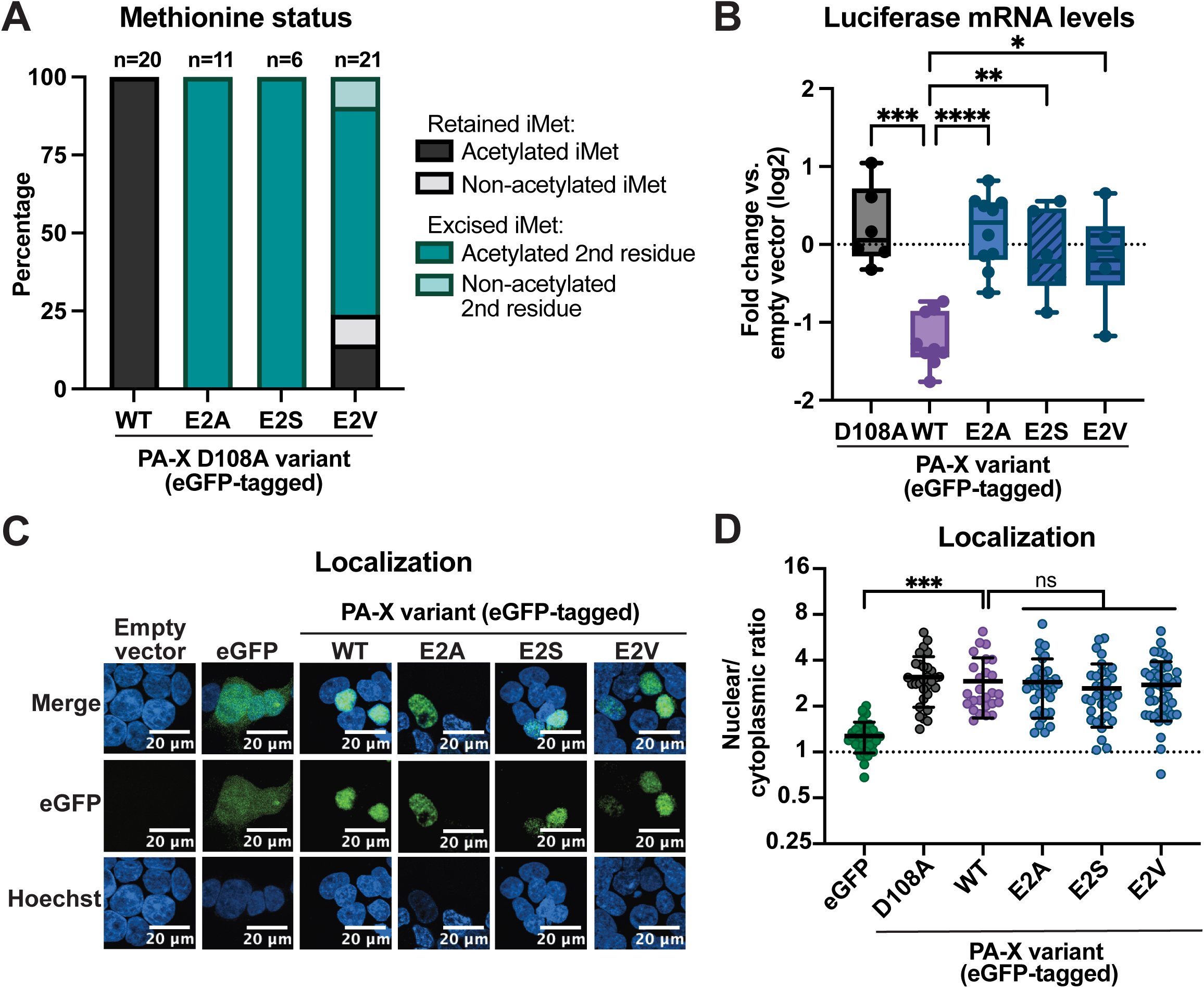
General N-terminal acetylation is sufficient for PA-X localization to the nucleus. HEK293T cells were transfected for 24 hours with empty vector, WT PR8 PA-X-eGFP, the catalytically inactive PA-X D108A-eGFP mutant, or the indicated NatA-modified mutants carrying changes in the second amino acid in the WT or D108A PR8-PA-X-eGFP background. A) eGFP-tagged proteins were isolated by immunoprecipitation and peptides were analyzed by nanoLC-MS/MS for initiator methionine (iMet) retention and N-terminal acetylation. N above stacked bars indicate total detected peptides for each variant over two experiments. B) mRNA levels of a co-transfected luciferase reporter were measured by RT-qPCR and normalized to 18S rRNA. Levels are plotted as fold change relative to vector transfected cells in log2 scale. C) Confocal microscopy was used to image eGFP, PA-X-eGFP (WT), PA-X E2A-eGFP (E2A), PA-X E2S-eGFP (E2S), and PA-X E2V-eGFP (E2V). Nuclei were stained with Hoechst. D) Nuclear/cytoplasmic ratios were calculated by measuring the mean fluorescence intensity of the nuclear and cytoplasmic compartments using ImageJ. Hoechst staining was used as a guide to generate regions of interest for the nucleus. Autofluorescence was used as a guide to generate cytoplasmic regions of interest. Nuclear/cytoplasmic ratios represent the following localization: <1: protein is primarily cytoplasmic, ∼1: protein is diffuse throughout the cell, >1: protein is primarily nuclear. For all experiments, some conditions were carried out in parallel to experiments in other figures. Therefore, control conditions (empty vector, D108A, WT, and eGFP) are identical. N≥3 for all experiments except the MS analysis in panel A. For microscopy, representative images are shown. In nuclear/cytoplasmic ratio quantitation, each data point represents 10-15 eGFP positive cells. * = p < 0.05, ** = p < 0.01, *** = p < 0.001, **** = p < 0.0001, ANOVA with Dunnett’s multiple comparison test.

### N-terminal acetylation at the initiator methionine by any Nat complex promotes PA-X host shutoff activity

It is surprising that PA-X mutants that are acetylated and localize to the nucleus (Fig. 2A,C,D) are inactive. A previous study hypothesized that perhaps the NatB complex activity, rather than the modification, was important(9). However, NatA– and NatB-modified proteins also differ in the presence of the initiator methionine and thus the location of the modification (Fig. 1A). To test if the site of the modification was the determining factor for PA-X activity, we investigated the effect of modification by NatC, which, like NatB, also modifies proteins on the initiator methionine(10,13). We generated two mutants that should be modified by NatC(13, 15), PA-X E2M and PA-X E2L (Fig. 1A). Mass spectrometry analysis showed that the initiator methionine of PA-X E2M was retained in 80% of peptides (27 out of 34), and about 60% of those peptides (15 out of 27) were N-terminally acetylated (Fig. 3A). For PA-X E2L, the initiator methionine was present on all detected peptides but only 20% (3 out of 14) were N-terminally acetylated (Fig. 3A). The subcellular localization of the NatC-modified mutants was similar to that of WT PA-X (Fig. 3B,C). In addition, the NatC-modified mutants were also able to downregulate luciferase mRNA levels in cells, although to a lesser degree than WT PA-X and the other NatB-modified mutants (Fig. 3D). The lower host shutoff activity may be linked to the incomplete acetylation of the proteins. Nonetheless, these results suggest that N-terminal acetylation of the initiator methionine specifically, and not NatB activity, is required for the host shutoff activity of PA-X.

**Fig. 3.**
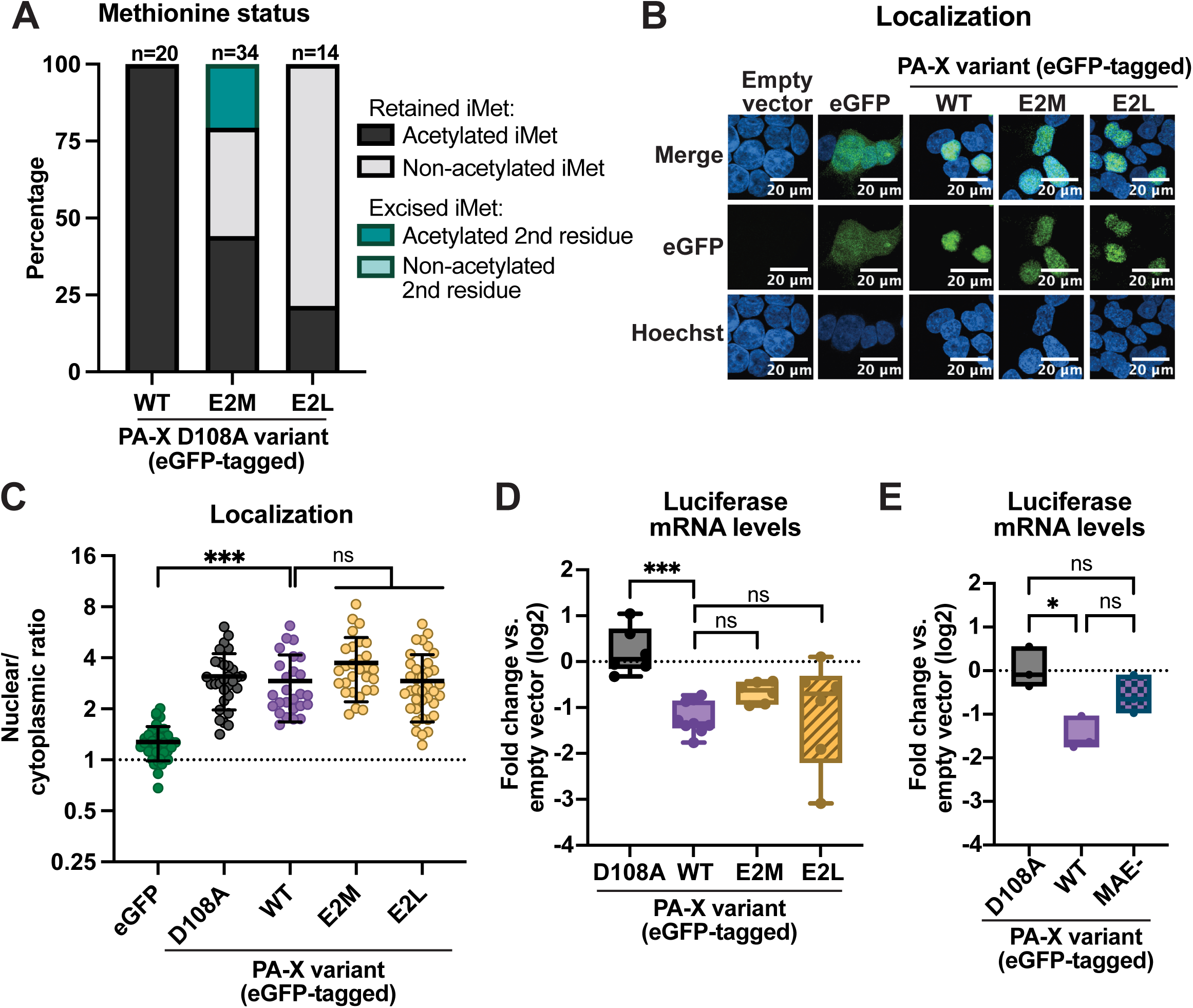
N-terminal acetylation at the initiator methionine promotes PA-X host shutoff activity. HEK293T cells were transfected for 24 hours with empty vector, WT PR8 PA-X-eGFP, the catalytically inactive PA-X D108A-eGFP mutant, the indicated NatC-modified mutants carrying changes in the second amino acid in the WT or D108A PR8-PA-X-eGFP background, or the NatA-modified PR8 PA-X MAE-eGFP mutant, which has an alanine insertion between the native first and second amino acid. A) eGFP-tagged proteins were isolated by immunoprecipitation and peptides were analyzed by nanoLC-MS/MS for initiator methionine (iMet) retention and N-terminal acetylation. N above stacked bars indicate total detected peptides for each variant over two experiments. B) Confocal microscopy was used to image eGFP, PA-X-eGFP ( WT), PA-X E2M-eGFP (E2M) and PA-X E2L-eGFP (E2L). Nuclei were stained with Hoechst. C) Nuclear/cytoplasmic ratios were calculated by measuring the mean fluorescence intensity of the nuclear and cytoplasmic compartments using ImageJ. Hoechst staining was used as a guide to generate regions of interest for the nucleus. Autofluorescence was used as a guide to generate cytoplasmic regions of interest. Nuclear/cytoplasmic ratios represent the following localization: <1: protein is primarily cytoplasmic, ∼1: protein is diffuse throughout the cell, >1: protein is primarily nuclear. D, E) mRNA levels of a co-transfected luciferase reporter were measured by RT-qPCR and normalized to 18S rRNA. Levels are plotted as fold change relative to vector transfected cells in log2 scale. For all experiments, some conditions were carried out in parallel to experiments in other figures. Therefore, control conditions (empty vector, D108A, WT, and eGFP) are identical. N≥3 for all experiments except the MS analysis in panel A. For microscopy, representative images are shown. In nuclear/cytoplasmic ratio quantitation, each data point represents 10-15 eGFP positive cells. ns = p > 0.05, * = p < 0.05, *** = p < 0.001, ANOVA with Dunnett’s multiple comparison test.

While these results show that the difference between the NatA and NatB-modified proteins is not the complex that deposits the modification, there are still two differences between proteins modified by the two complexes – the removal of one amino acid in ME/D/N-vs. (M)A-starting proteins and the specific residue that is modified (M vs. A). Both features have the potential to change the surface of the protein, which in turn may alter PA-X interactions with RNA or proteins. To distinguish the effects of these two differences, we generated a mutant that should have its N-terminal methionine excised while remaining the same length as WT PA-X and bearing a glutamic acid at the second position. We achieved this by inserting an additional alanine between the initiator methionine and the second residue, generating the PA-X MAE-mutant. The PA-X MAE-mutant should lose its initiator methionine and be modified on the alanine by NatA. Thus, the only difference between WT PA-X and PA-X MAE-should be that the acetylation occurs on an alanine instead of a methionine. Interestingly, PA-X MAE-still largely lost the ability to downregulate luciferase mRNA levels (Fig. 3E). These results indicate that the initiator methionine itself is also important.

### Neither eGFP tagging nor expression levels of the E2 mutant proteins explain the differences in localization or activity

Although the host shutoff activity of PA-X-eGFP was similar to that of untagged PA-X (Fig. 4A), we explored the possibility that our detection strategy for PA-X may lead to unexpected confounds due to the size of the eGFP tag. This is especially relevant for protein localization. We therefore tested whether a smaller C-terminal 3xFlag tag would yield the same results. Like PA-X-eGFP, WT PA-X-3xFlag had similar host shutoff activity to untagged PA-X (Fig. 4A). We then generated PA-X E2A, PA-X E2P, PA-X E2D, and PA-X E2M 3xFlag-tagged proteins, and found that they had similar host shutoff activity to their eGFP-tagged counterparts, corroborating our findings on the role of N-terminal acetylation on the initiator methionine for activity (Fig. 4B). In immunofluorescence assays, WT PA-X-3xFlag and all the PA-X E2 mutant-3xFlag-tagged proteins had a more diffuse localization than their eGFP-tagged counterparts (Fig. 4C). While this could be due to the tag difference, we note that we transfected cells with 16X the amount of PA-X-3xFlag containing plasmids to ensure detection, which may increase the overall levels of PA-X and thus the portion of PA-X found outside the nucleus. Nonetheless, the E2 mutations altered Flag-tagged PA-X localization in an analogous manner to the eGFP-tagged PA-X. WT PA-X, PA-X E2D (both NatB-modified), and PA-X E2M (NatC-modified) had similar nuclear/cytoplasmic ratios, approximately 0.8, 1.4, and 0.8 respectively. PA-X E2A (NatA-modified) was slightly less nuclear (ratio ∼0.5), although this difference did not reach statistical significance when compared to WT PA-X (Fig. 4D). In contrast, the unmodified PA-X E2P-3xFlag was significantly less nuclear than WT PA-X (ratio ∼0.3, Fig. 4D). Overall, the similar trends between the eGFP-tagged and 3xFlag-tagged PA-X E2 mutants support the conclusion that N-terminal acetylation at any position is sufficient for PA-X nuclear localization.

**Fig. 4.**
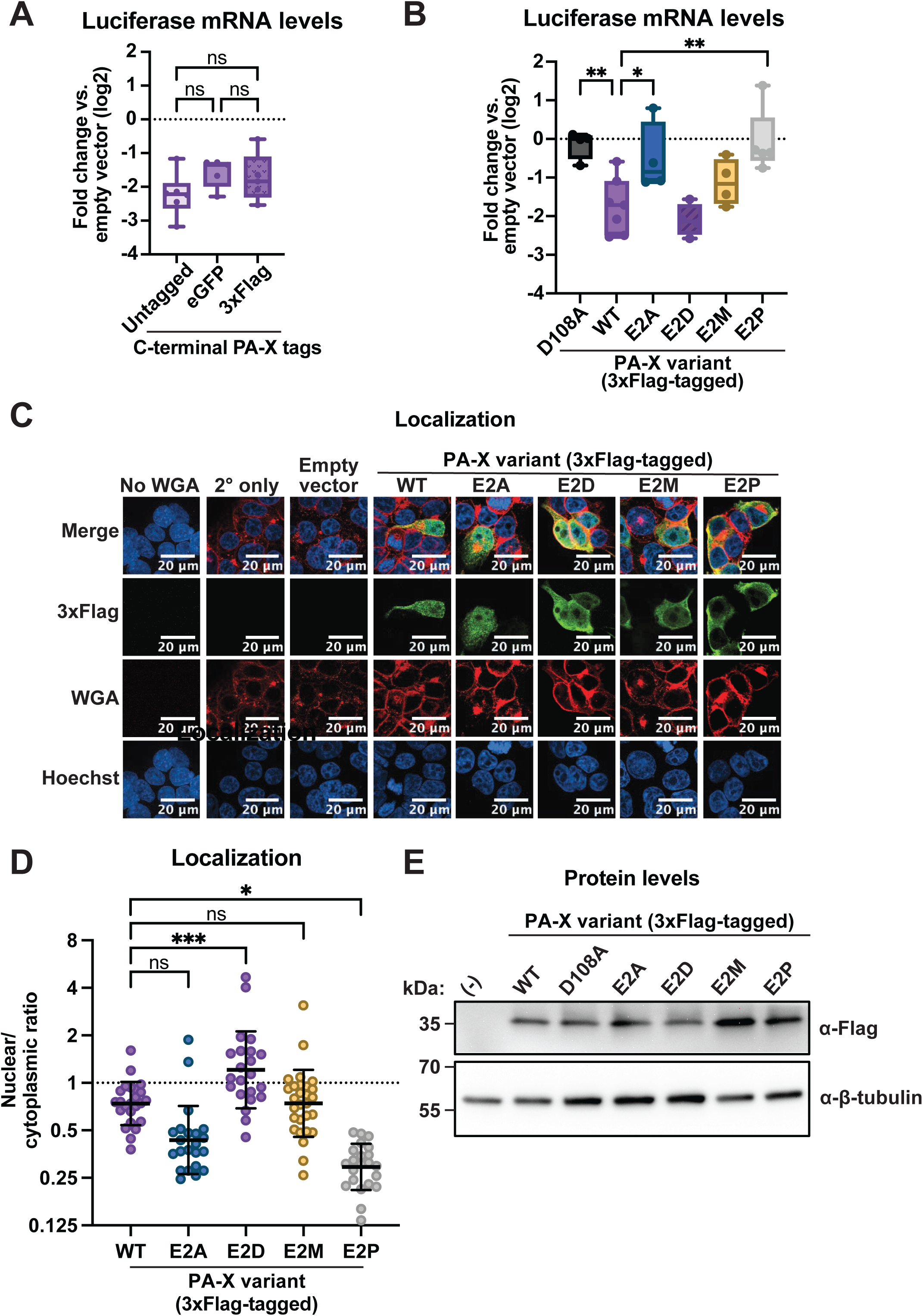
Differences in PA-X localization and activity are not due to the eGFP tag or variations in protein levels. HEK293T cells were transfected for 24 hours with empty vector, untagged WT PR8 PA-X, PA-X-eGFP, PA-X-3xFlag, or the following 3xFlag-tagged PA-X mutants: E2D, E2A, E2M, or E2P. A, B) mRNA levels of a co-transfected luciferase reporter were measured by RT-qPCR and normalized to 18S rRNA. Levels are plotted as fold change relative to vector transfected cells in log2 scale. C) Confocal microscopy was used to image PA-X-3xFlag (WT), PA-X E2D-3xFlag (E2D), PA-X E2A-3xFlag (E2A), PA-X E2M-3xFlag (E2M), and PA-X E2P-eGFP (E2P). Cell membranes were stained with WGA (red), PA-X was stained with α-Flag antibodies (green) and nuclei were stained with Hoechst (blue). D) Nuclear/cytoplasmic ratios were calculated by measuring the mean fluorescence intensity of the nuclear and cytoplasmic compartments using ImageJ. Hoechst and WGA staining were used as guides to generate regions of interest for the nucleus and cell membrane, respectively. E) Protein levels were analyzed by western blotting. A Flag antibody was used to visualize Flag-tagged PA-X and PA-X mutants. Tubulin staining was included as a loading control. (−) denotes empty vector transfection. N≥3 for all experiments. For microscopy and western blots, representative images are shown. In nuclear/cytoplasmic ratio quantitation, each point represents 1-3 PA-X-positive cells. ns = p > 0.05, *p < 0.05, ***p < 0.001, ANOVA with Dunnett’s multiple comparison test.

We have also struggled to detect the eGFP-tagged PA-X proteins consistently by western blotting, which limited our ability to ensure that changes to the PA-X sequence did not alter its levels. We thus tested whether the 3xFlag tag, which has been used successfully by others(7), gave us better detection. Indeed, we found that we could easily visualize Flag-tagged PA-X on western blot. Moreover, we were able to confirm that WT and the E2 mutants were expressed at similar levels, including the inactive E2A and E2P mutants (Fig 4E). Therefore, we can conclude that expression differences do not explain activity or localization changes due to these mutations.

### N-terminal acetylation by different Nat complexes has similar consequences on the activity of PA-X of all influenza A strains tested

While results with WSN(9) and PR8 PA-X are suggestive of a broader phenomenon, both of these strains are lab-adapted viruses. Therefore, we sought to ensure that the phenotype of N-terminal acetylation mutants was conserved in PA-X proteins from more relevant human seasonal influenza viruses. Currently circulating seasonal human viruses are H1N1 strains related to the pandemic 2009 H1N1 virus (H1N1pdm09), and H3N2 strains(44, 45). Thus, we tested the effect of E2P (unmodified), E2A (NatA-modified), and E2M (NatC-modified) mutations in the PA-X sequences from A/Tennessee/1-560/2009 H1N1 (H1N1pdm09) and A/Perth/16/2009 H3N2 (Perth). We found that the trend in host shutoff activity was similar among PA-X proteins from all influenza A strains tested. All PA-X E2A and PA-X E2P mutants had significantly lower (or no) host shutoff activity, while the PA-X E2M mutants maintained an intermediate amount of activity (Fig. 5). These results suggest that regardless of viral strain, PA-X requires N-terminal acetylation in general to access the nucleus and N-terminal acetylation of the initiator methionine specifically to downregulate RNA levels.

**Fig. 5.**
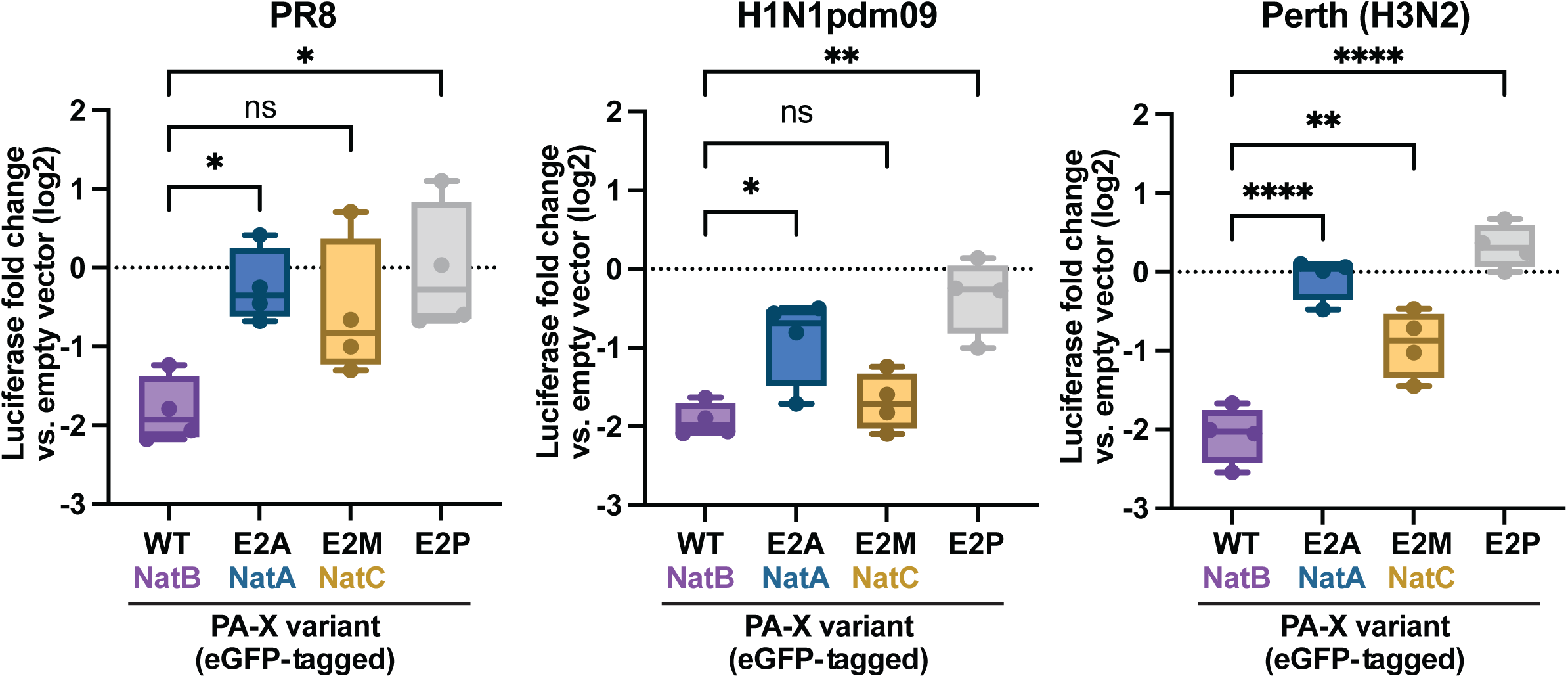
N-terminal acetylation also regulates the activity of PA-X proteins from circulating influenza A virus subtypes. HEK293T cells were transfected for 24 hours with empty vector, WT PR8 PA-X-eGFP, WT H1N1pdm09 PA-X-eGFP, or WT Perth PA-X-eGFP or the corresponding PA-X-eGFP mutants carrying changes in the second amino acid. mRNA levels of a co-transfected luciferase reporter were measured by RT-qPCR and normalized to 18S rRNA. Levels are plotted as fold change relative to vector transfected cells in log2 scale. n=4. ns = p > 0.05, * = p < 0.05, ** = p < 0.01, ****= p < 0.0001, ANOVA with Dunnett’s multiple comparison test.

### N-terminal acetylation is required for PA-X host shutoff activity during infection

Our results point to the importance of PA-X N-terminal acetylation for nuclear localization and host shutoff activity when PA-X is ectopically expressed in cells. However, we wanted to confirm that this modification is also important during viral infection. This was not done in the previous study by Oishi et al.(9) because of the challenges in separating PA and PA-X modifications during infection. PA-X is produced by a +1 frameshifting event that occurs after translation of the 191st amino acid of PA(1, 35). Therefore, PA and PA-X have the same N terminus and any mutation in the PA-X N terminus is also present in PA(1, 35). This is problematic as PA also requires N-terminal acetylation for polymerase activity(9). Thus, to separate PA and PA-X production, we examined the activity of ectopically expressed PA-X mutants in cells infected with PA-X-deficient PR8 (PR8 PA(ΔX))(5). While the human lung epithelial cell line A549 is most commonly used as a model for influenza infection, transfection efficiency is low in these cells.

Therefore, we used HEK 293T cells, which are readily transfectable and have been used in many previous studies of influenza A virus infections(46–54). To ensure that the experimental setup would work, we first checked for potential interference between transfection and infection. We transfected HEK293T cells with plasmids encoding eGFP or eGFP-tagged WT PA-X, left them uninfected or infected them with WT PR8 or PR8 PA(ΔX), and tested for the efficiency of PA-X host shutoff and of infection. We found that transfecting cells did not prevent them from being infected with WT PR8 or PR8 PA(ΔX) viruses, as all cells expressed similar levels of influenza HA RNAs (Fig. 6A). Moreover, transfection did not inhibit the ability of virus-encoded PA-X to downregulate endogenous targets, as G6PD, a known target of PA-X(5), was downregulated in transfected cells that were infected with WT PR8, but not PR8 PA(ΔX) (Fig. 6B). We then repeated the experiment and transfected cells with a luciferase reporter and eGFP-tagged versions of WT PA-X, PA-X E2D, which is still modified by NatB, PA-X E2P, which is unmodified, or eGFP alone. Cells were then infected with PR8 PA(ΔX) virus or mock infected (Fig. 6C). As expected, WT PA-X was active and downregulated luciferase mRNA levels in both mock and PR8 PA(ΔX)-infected cells and the E2D mutant had similar activity as WT PA-X under both conditions. In contrast, based on luciferase mRNA downregulation, unmodified PA-X E2P was inactive in both mock and PR8 PA(ΔX)-infected cells. This result confirms that PA-X also requires N-terminal acetylation for activity during infection.

**Fig. 6.**
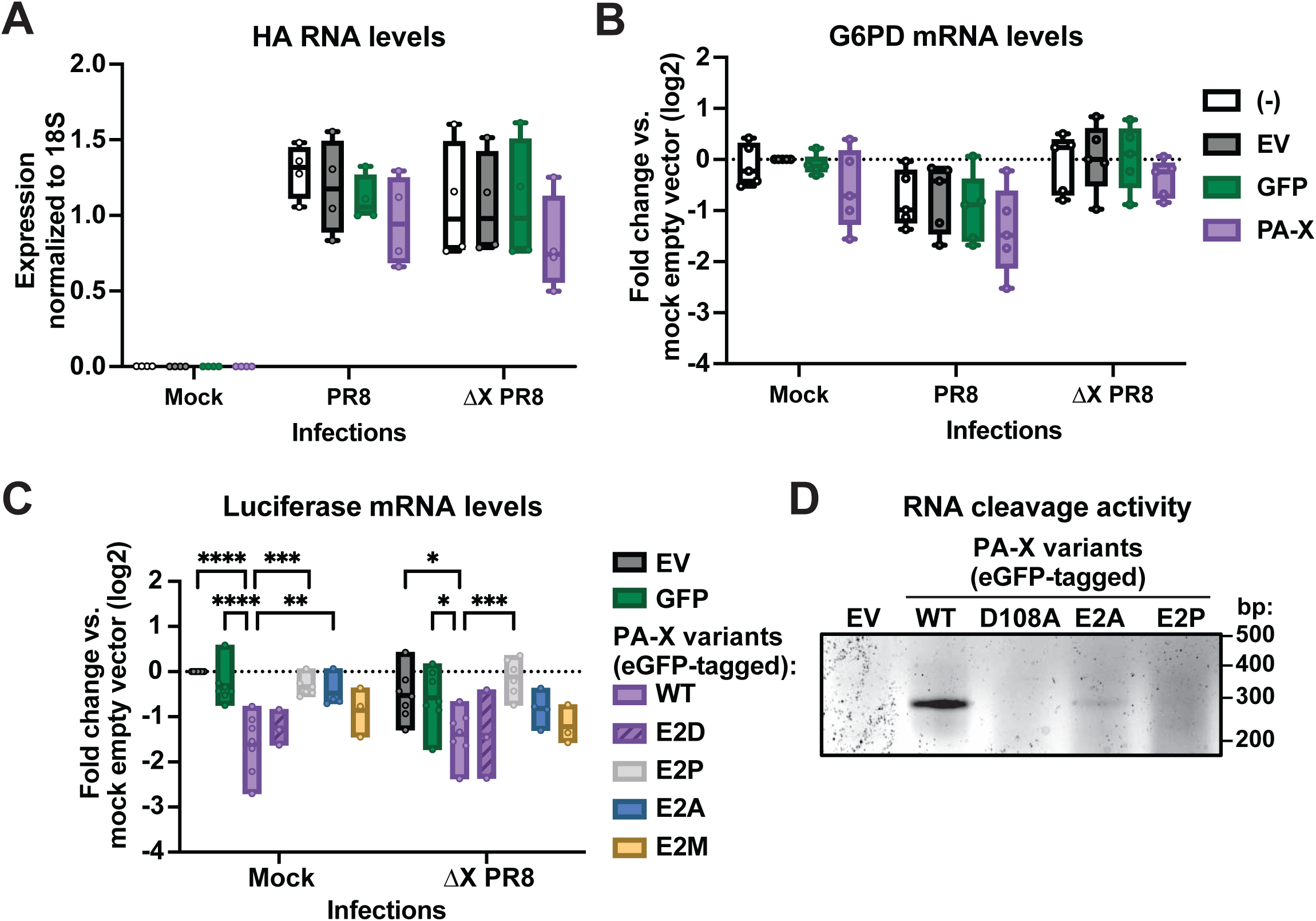
During infection, N-terminal acetylation of PA-X at any position is sufficient for host shutoff activity. A-C) HEK293T cells were transfected for 24 hours with empty vector (EV), eGFP, and eGFP-tagged WT PA-X or the following eGFP-tagged mutants: unmodified mutant PA-X E2P (gray), NatB-modified PA-X E2D mutant (purple), NatA-modified mutant E2A (blue), or NatC-modified mutant E2M (yellow). Eight hours post-transfection, cells were mock infected or infected with WT PR8 or PA-X deficient PR8 (ΔX) at an MOI of 1. RNA samples were collected 16 hrs post-infection (24 hrs post-transfection). RNA levels of A) viral HA RNA, B) endogenous G6PD mRNA, and C) the co-transfected luciferase reporter were measured by RT-qPCR and normalized to 18S rRNA. For B,C, levels are plotted as change relative to vector transfected cells in log2 scale. n≥3. * = p < 0.05, ** = p < 0.01, *** = p < 0.001, **** = p < 0.0001, Ordinary two-way ANOVA with uncorrected Fisher’s LSD multiple comparison test compared to the wild-type PA-X in each infection condition. D) HEK293T ishXRN1 cells were transfected for 24 hours with empty vector and the following eGFP-tagged constructs: WT PA-X, PA-X D108A (catalytically inactive), PA-X E2A, or PA-X E2P, alongside the pCMV-luciferase+intron-YKT6-99bp cleavage reporter. RNA samples were collected 24 hours post-transfection and 5’ RACE was performed. Bands seen on the agarose gel correspond to the predicted size of the PCR product corresponding to the cleaved fragment (290 bp) and its provenance was verified by Sanger sequencing. The agarose gel image is representative of 3 biological experiments.

### During infection, N-terminal acetylation at any position is sufficient for PA-X host shutoff

To test whether other requirements for PA-X activity were also similar during infection, we transfected cells with PA-X carrying E2A (NatA-modified), or E2M (NatC-modified) mutations. To our surprise, NatA-modified PA-X E2A had higher levels of host shutoff activity in PR8 PA(ΔX)-infected vs. mock-infected cells (Fig. 6C). As in other experiments using ectopic expression (Figs. 1-5), luciferase downregulation by PA-X E2A was significantly reduced compared to downregulation by WT PA-X in mock infected cells. In contrast, during infection, luciferase downregulation by PA-X E2A was not significantly different from downregulation by WT PA-X. Luciferase downregulation by PA-X E2M was not significantly different from that by WT PA-X in mock infected cells, consistent with our previous assays (Figs. 3-5), nor in infected cells (Fig. 6C).

The discrepancy between the effects of PA-X E2A on steady state RNA levels in uninfected vs. infected cells suggests that this mutant retains RNase activity, even though it has minimal impact on steady state mRNA levels in uninfected cells. To confirm this, we used 5’ RACE to test its cleavage activity on a known PA-X target sequence from the human YKT6 mRNA inserted into the luciferase reporter, an assay we have previously used to study PA-X targeting preferences(6). Since 5’ RACE is an end-point PCR-based assay, it does not accurately reveal quantitative difference in protein activity, but provides only a binary cleavage/no cleavage readout. Interestingly, even though ectopically expressed PA-X E2A did not downregulate luciferase mRNA by RT-qPCR (Fig. 2B), it was able to cleave mRNA targets at the previously identified PA-X target site to some extent (Fig. 6D). This suggests that PA-X E2A mutants retain RNase activity but somehow cannot properly reach their targets in uninfected cells. An additional factor must increase the ability of PA-X E2A to cleave RNA in the context of infection, perhaps by promoting its association with the mRNA targets. Taken together, these results suggest that additional factors during infection potentiate PA-X host shutoff activity, a phenomenon that is only apparent in the context of PA-X mutants with attenuated host shutoff activity.

### The increase in PA-X activity during infection cannot be explained by any single influenza A virus protein

A possible reason for the increased host shutoff activity of PA-X E2A during infection is a functional interaction between PA-X and another viral protein. In the past, this possibility has not been heavily investigated, since all studies have indicated that PA-X is active upon ectopic expression in uninfected cells, in the absence of other viral proteins. Also, our previous analyses suggest that PA-X has similar RNA targets in infected and uninfected cells(4–6). However, one recent study indicated that NS1 has a role in PA-X-mediated host shutoff(55). To test the hypothesis that other influenza proteins potentiate PA-X activity, we compared mRNA downregulation by ectopically expressed WT PA-X alone or by the NatA-modified PA-X E2A mutants expressed alongside individual influenza proteins from the PR8 strain (Fig. 7A). To prevent WT PA-X expression from the PA segment, we used a modified construct with a mutation in the frameshift sequence (TTTCGT to TTCCGC) that strongly attenuates the +1 ribosomal frameshift during translation and reduces PA-X production(56). We were unable to express PB2, PB1, or HA at sufficient levels (Fig. 7A). Of the co-expressed proteins that we detected by western blot, only NS1 appeared to increase the activity of PR8 E2A (Fig. 7B,C), However, upon subsequent experiments, we found that PR8 NS1 alone caused a small downregulation in luciferase RNA in our experiments (Fig. 7C,D). NS1-PA-X E2A coexpression increased downregulation of co-expressed luciferase mRNA and GFP protein relative to PA-X E2A alone, but did not completely restore wild-type levels of PA-X activity (Fig. 7C,D). Therefore, the increase in activity could simply be due to an additive effect between PA-X E2A and NS1. Collectively, these results suggest that none of the tested influenza A virus proteins can explain the potentiated activity of PA-X during influenza A virus infection.

**Fig. 7.**
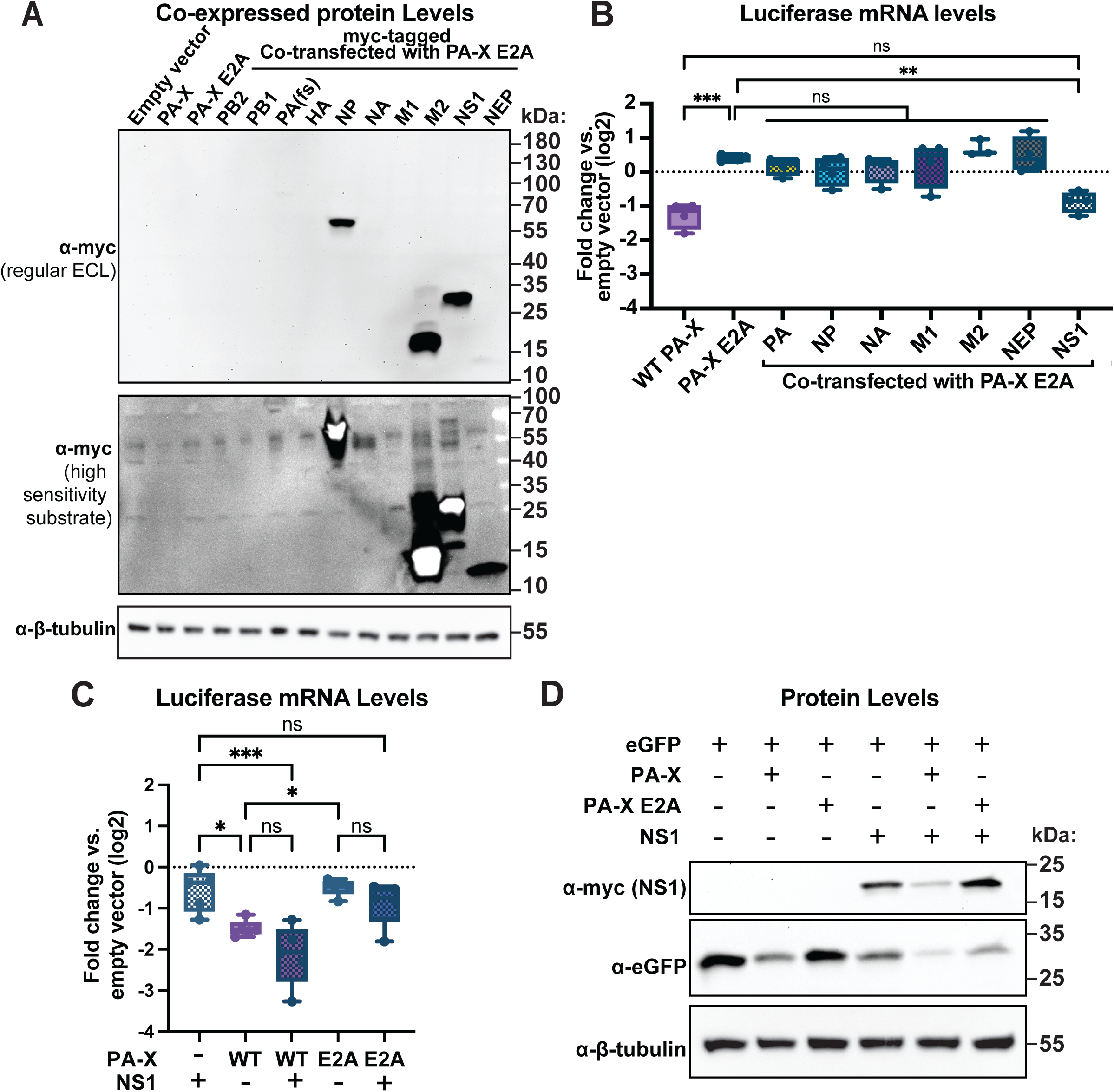
The increase in PA-X activity during infection is not due to the action of any single other influenza A virus protein. A,B) HEK293T cells were transfected for 24 hours with empty vector, WT PR8 PA-X-eGFP, the NatA-modified mutant PA-X E2A-eGFP, or PA-X E2A-eGFP together with myc-tagged versions of the indicated PR8 proteins. A) Production of the PR8 proteins was analyzed by western blotting using an antibody against the myc tag, with tubulin as a loading control. Chemiluminescent signal was generated using the Pierce ECL Western Blotting Substrate (regular ECL) or SuperSignal West Atto Ultimate Sensitivity Substrate (high sensitivity substrate). B) mRNA levels of a co-transfected luciferase reporter were measured by RT-qPCR and normalized to 18S rRNA. Levels are plotted as fold change relative to vector transfected cells in log2 scale. C,D) HEK293T cells were transfected for 24 hours with empty vector, eGFP, WT PA-X-eGFP, the NatA-modified mutant PA-X E2A-eGFP, PR8 NS1-myc or a combination of these constructs, as indicated. C) mRNA levels of a co-transfected luciferase reporter were measured by RT-qPCR and normalized to 18S rRNA. Levels are plotted as fold change relative to vector transfected cells in log2 scale. D) Protein levels were analyzed by western blotting. A myc antibody was used to visualize NS1 and a GFP antibody to visualize eGFP. Tubulin staining was included as a loading control. eGFP-tagged PA-X was not detected in these lysates, despite clear mRNA downregulation in C, likely due to low levels of expression. This is consistent with what we have observed in other experiments. N≥3 for all experiments. For western blots, representative images are shown. ns = p > 0.05, *p < 0.05, **p < 0.01, ***p < 0.001, ANOVA with Dunnett’s multiple comparison test.

## Discussion

Our results reveal that N-terminal acetylation of PA-X supports host shutoff activity in two separate ways: it is required for nuclear localization of PA-X, and it is involved in a second yet undetermined process that promotes PA-X activity. Interestingly, the two processes have slightly different requirements for the modification. Addition of the acetyl group at either the first residue, i.e. the initiator methionine, or the second residue following initiator methionine excision both promote nuclear localization of PA-X (Fig. 1D,E, 2C,D). However, mutants that are modified at the second residue, like PA-X E2A, still have reduced host shutoff activity and do not strongly downregulate RNA levels (Fig. 2B), despite being localized to the nucleus (Fig. 2C,D) and retaining RNase activity (Fig. 6D). Further mutational analysis indicates that the methionine residue at the beginning of the protein is important, rather than the change in protein size and potentially protein surface (Fig. 3E). Therefore, our results indicate that an acetyl group on the initiator methionine is specifically required for the full host shutoff activity of PA-X, at least when expressed ectopically. Interestingly, we found that the retained methionine is not required for the host shutoff activity of PA-X during infection (Fig. 6C). This restoration of host shutoff activity during infection may be due to a combination of the other viral proteins and/or cellular factors that change during infection.

How N-terminal acetylation separately mediates the nuclear localization and the host shutoff activity of PA-X remains unclear, although these effects are likely linked to how this modification impacts protein-protein interactions(25–28). Biochemically, N-terminal acetylation eliminates the N-terminal positive charge of the first amino acid and increases hydrophobicity. This may allow for a new protein interaction surface that is unavailable when the N-terminus is unmodified. For example, the N-terminal acetylation of initiator methionine in the NEDD8-conjugating E2 enzyme Ubc12 allows for its docking into the hydrophobic pocket of the co-E3 ligase Dcn1, thus promoting the function of the E2/E3 complex(25, 26). The structure of the interaction interfaces may also be subtly different depending on whether the initiator methionine is retained. In the case of PA-X, N-terminal acetylation at either the first or second amino acid may both support interactions with a cytoplasmic protein that is involved in the nuclear import of PA-X. However, retention of a modified methionine may be needed to create the correct interaction surface for a protein that allows PA-X to reach, bind, and/or degrade its RNA targets in the nucleus. These possibilities should be addressed in future studies.

This is the first time that a role for the N-terminal region of PA-X in nuclear localization has been reported. Previous studies have identified charged residues in the C-terminal X-ORF of PA-X that are important for nuclear localization(4, 7, 36). Here we show that these sequences alone are not enough for PA-X nuclear localization, as they were not mutated in our studies. Conversely, N-terminal acetylation is also not enough to direct PA-X nuclear localization, because the N-terminal domain of PA-X alone does not traffic to the nucleus(4, 7, 36), even though it is likely still N-terminally acetylated by NatB. The full structure of PA-X with its C-terminal X-ORF has not yet been solved, and further work should determine how the acetylated N-terminus and the C-terminal X-ORF work together to drive the nuclear localization of PA-X. Our study is also the first report of regulation of nuclear localization by N-terminal acetylation. Other post-translational modifications like phosphorylation(57, 58), arginine methylation(59–61), and lysine acetylation(62) are well-known to modulate nuclear localization, largely by mediating interactions with importin α, which mediates nuclear import of proteins. In contrast, N-terminal acetylation has only previously been described to regulate protein localization to membranes. N-terminal acetylation directs proteins to membranes through protein-protein interactions with integral membrane proteins(32, 33) or direct interactions with membranes(63). It is possible that the N-terminal acetylation similarly increases binding affinity between PA-X and interacting proteins that mediate nuclear localization. While the details of this effect are still unknown, these results suggest that N-terminal acetylation has a broader range of functions in protein localization than previously reported and mediates interactions with additional trafficking proteins.

The glutamic acid at the second position of PA-X is highly conserved among influenza isolates, with 99.7% of H1N1pdm09 viruses, 99.9% of H3N2 viruses, 100% of H5N1, and 100% of H7N9 viruses containing this amino acid at the second position(9). This high conservation underscores how N-terminal acetylation of PA-X by NatB is important for the host shutoff activity of PA-X. We were thus surprised to find that during infection, the NatA-modified PA-X E2A mutant has almost wild-type activity, even though PA-X E2A alone does not downregulate mRNA levels and alanine at the second position of PA-X only rarely occurs. Because PA-X E2A retains RNA cleavage activity during ectopic expression, we conclude that this mutant is partially active and that we can use it to reveal additional factors during infection that support PA-X activity. We investigated other viral proteins first, as a study by Bougon et al. found some NS1 mutations abolished host shutoff in PR8-infected cells, pointing to a functional interaction between PA-X and NS1(55). However, in our experiments the effect of NS1 addition is small and does not explain the difference between PA-X E2A activity during ectopic expression vs. infection. Because influenza A virus infection alters the levels of some host proteins(64, 65), changes in cellular proteins may also promote PA-X host shutoff activity. In particular, proteins with functions in protein localization and transport are upregulated in influenza A virus-infected human lung cells(64). Higher levels of a PA-X interacting protein could compensate for a reduction in PA-X binding affinity resulting from changes in N-terminal acetylation in the PA-X E2A mutant. In general, our results point to additional contributions in the regulation of PA-X activity, both NS1-dependent and independent, which will need to be defined in the future.

Overall, we have discovered that N-terminal acetylation functionally promotes the host shutoff activity of PA-X by separately influencing PA-X subcellular localization and its overall ability to downregulate target mRNAs. Our results also indicate that acetylation of the initiator methionine specifically promotes RNA downregulation by PA-X. In the future, it will be important to determine how N-terminal acetylation contributes to structural differences and/or PA-X interactions with other proteins and PA-X substrates. Additionally, in the context of cellular biology, these studies contribute new evidence that N-terminal acetylation supports the correct subcellular localization of proteins and, to our knowledge, constitute the first report that N-terminal acetylation promotes nuclear localization. Our studies also highlight how N-terminal acetylation is multifaceted in its regulations of modified proteins and how this modification remains incompletely understood, limiting our ability to discern its role in the regulation of proteins in physiological and pathological settings.

## Materials and Methods

### Plasmids

C-terminally tagged pCR3.1-PA-X-eGFP, pCR3.1-PA-X D108A-eGFP, pCR3.1-PB2-myc, pCR3.1-PB1-myc, pCR3.1-PA(fs)-myc, pCR3.1-HA-myc, pCR3.1-NP-myc, pCR3.1 NA-myc, pCR3.1-M1-myc, pCR3.1-NS1-myc, and pCR3.1-NEP-myc from the PR8 strain were gifts from C. McCormick (Dalhousie University, Halifax, NS, Canada) and generated as previously described(56). pCDNA3.1-eGFP and pCDNA4/TO-C-terminal 3xFlag were a gift from B. Glaunsinger (University of California, Berkeley, Berkeley, CA, USA). The luciferase construct with the β-globin intron (pCMV luciferase + intron) was a gift from G. Dreyfuss (University of Pennsylvania, Philadelphia, PA, USA)(66). The pCMV-luciferase+intron-YKT6-99bp construct containing a strong PA-X cut site from the human YKT6 gene within the luciferase mRNA was previously described(6). The rescue plasmids encoding the 8 segments of PR8 virus (pHW191-PB2 to pHW198-NS) were gifts from R. Webby (St. Jude Children’s Research Hospital, Memphis, TN, USA)(67). Gibson assembly using the HiFi assembly mix (New England Biolabs) was used to make pCR3.1-PA-X-NLS-eGFP by amplifying the PR8 sequence of PA-X from pCR3.1-PA-X-myc and the eGFP sequence from pCDNA3.1-eGFP. The nuclear localization sequence (NLS) in pCR3.1-PA-X-NLS-eGFP was designed into the primers for Gibson assembly. From this construct, we then generated the PA-X E2P-NLS mutant using QuikChange II site-directed mutagenesis (Agilent). eGFP fusion constructs for PA-X from non-PR8 strains were made by replacing the PR8 PA-X sequence in pCR3.1-PA-X-eGFP with the PA-Xs from A/Perth/16/2009 H3N2 (Perth) and A/Tennessee/1-560/09 H1N1 (H1N1pdm09) subcloned from previously published myc-tagged constructs(4, 6) using the MluI/SalI restriction sites and T4 DNA ligation (New England Biolabs). The E2 mutations and the MAA– and MAE-insertions in the pCR3.1-PA-X-eGFP constructs with PR8 PA-X, PR8 PA-X D108A, Perth PA-X, and H1N1pdm09 PA-X were generated using QuikChange II site-directed mutagenesis (Agilent). The C-terminal 3xFlag-tagged PA-X constructs were generated by amplifying WT and E2 mutant PR8 PA-X sequences from the pCR3.1 backbones and inserting them into the HindIII and NotI sites of pCDNA4/TO-C terminal 3xFlag vector using Gibson cloning (HiFi assembly mix, New England Biolabs). The PR8 pHW-PA(ΔX) plasmid, previously described(5, 6), was generated from pHW193 by introducing mutations that reduce frameshifting events and add a premature stop codon in the PA-X reading frame, but that are silent in the PA reading frame.

### Cell lines and transfections

Human embryonic kidney (HEK) 293T cells and Madin-Darby canine kidney (MDCK) cells were commercially obtained from ATCC (CRL-3216 and CCL-34, respectively). HEK293T cells expressing inducible short hairpin RNA against XRN1 (ishXRN1) were previously described(68). All cell lines were maintained in high glucose Dulbecco’s modified Eagle’s medium (DMEM, Gibco) supplemented with 10% fetal bovine serum (FBS, Cytiva) at 37°C in 5% CO2 atmosphere. To measure host shutoff activity during transfection, HEK293T cells were plated on 24-well plates and transfected with 500 ng total DNA (including 25 ng PA-X constructs and 50 ng pCMV luciferase + intron construct) in 10% FBS in DMEM using jetPRIME transfection reagent (Polyplus). Cells were collected 24 hours post-transfection for RNA extraction and purification. To determine subcellular localization of PA-X using eGFP-tagged proteins, HEK293T cells were transfected as described above. To determine subcellular localization of PA-X using Flag-tagged proteins, HEK293T cells in 12-well plates were transfected with 500 ng of PA_X constructs. In both cases, they were plated onto poly-L-lysine (Sigma)-treated glass coverslips. To measure direct cleavage of RNAs by 5’ rapid amplification of DNA ends (RACE), HEK293T ishXRN1 cells were treated with 1 μg/mL doxycycline for 3-4 days to induce the XRN1 shRNA and knock-down the protein. They were then plated in 6-well plates and transfected with 2 μg total DNA, including 125 ng of PA-X constructs and 250 ng of the pCMV-luciferase+intron-YKT6-99bp cleavage reporter, similarly to what was done in Gaucherand et al.(6). To measure the host shutoff activity of PA-X during influenza A virus infections, HEK293Ts were plated in 24-well plates that were pre-treated with fibronectin, bovine serum albumin (BSA) and bovine collagen to increase HEK293T adhesion to the plate during the procedure. For the plate treatment, 10 ug/ml fibronectin (Sigma-Aldrich), 100 ug/ml BSA (Sigma-Aldrich), and 30 ug/ml bovine collagen (Advanced Biomatrix) in modified Eagle’s medium (MEM, Gibco) were added to the wells and UV crosslinked for 30 minutes. Wells were then washed three times with Dulbecco’s phosphate buffered saline (DPBS, Gibco) before addition of the cells. After plating, cells were transfected with 500 ng total DNA (including 25 ng PA-X constructs and 50 ng pCMV luciferase + intron construct) in infection media (0.5% low-endotoxin BSA (Sigma-Aldrich) in high glucose DMEM) using jetPRIME transfection reagent. 8 hours post-transfection, media was removed, and cells were infected as described below. 24 hours post-transfection/16 hours post-infection, cells were collected for RNA extraction and purification.

### Viruses and infections

Wild-type influenza A virus A/Puerto Rico/9/1934 H1N1 (PR8) and the mutant recombinant virus PR8 PA(ΔX) were generated using the 8-plasmid reverse genetic system(67) as previously described(4–6). Viral stocks were produced in MDCK cells and infectious titers were determined by plaque assays in MDCK cells using 1.2% Avicel (FMC BioPolymer) overlays(69). Briefly, confluent MDCK cells were infected with low volumes of tenfold serially diluted virus stocks in triplicate for 1 hour. Cells were then washed twice with DPBS before the addition of overlay media (1.2% Avicel, 1X MEM, 0.5% low-endotoxin BSA (Sigma-Aldrich), and 1 ug/ml TPCK-treated trypsin (Sigma-Aldrich)) and incubated for 4 days at 37°C in 5% CO_2_ atmosphere. After 4 days, cells were fixed with 4% paraformaldehyde and stained with crystal violet (Sigma-Aldrich) to observe plaques. Influenza A virus infections following HEK293T transfections were performed in infection media (0.5% low-endotoxin BSA in high glucose DMEM). Briefly, transfected HEK293T cells were treated with infection media supplemented with 0.5 ug/ml TPCK-treated trypsin alone (mock infection) or containing WT PR8 or PR8 PA(ΔX) at a multiplicity of infection (MOI) of 1 and incubated for 16 hours at 37°C in 5% CO_2_ atmosphere. Cells were then collected and lysed in RNA lysis buffer for RNA isolation.

### RNA purification, cDNA preparation and qPCR

RNA was extracted and purified using the Quick-RNA miniprep kit (Zymo Research) following the manufacturer’s protocol. Purified RNA was treated with Turbo DNase (Life Technologies), then reverse transcribed using iScript supermix (Bio-Rad) per manufacturer’s protocol. qPCR was performed using iTaq Universal SYBR Green Supermix (Bio-Rad) on the Bio-Rad CFX Connect Real-Time PCR Detection System or CFX Duet Real-Time PCR System and analyzed with Bio-Rad CFX Manager software or Bio-Rad CFX Maestro software. 18S ribosomal RNA was chosen as a housekeeping gene for normalizing data because its levels are stable during influenza A virus infections(70) and because rRNAs are not targeted by degradation by PA-X(4). Primers used for the qPCR were:

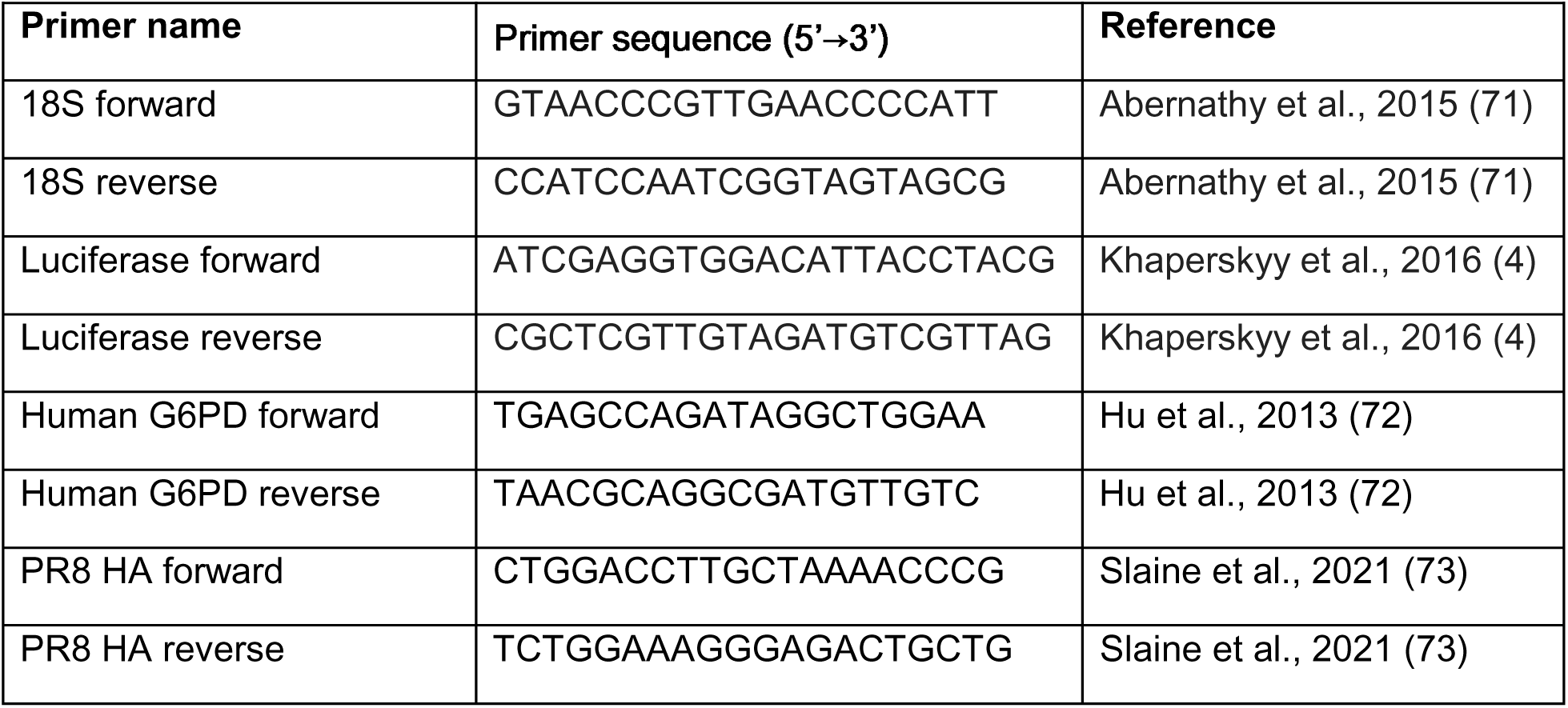

### Immunofluorescence assays and confocal microscopy

HEK293T cells were grown on glass coverslips pretreated with poly-L-lysine (Sigma-Aldrich), which were prepared following the manufacturer’s protocol. 24 hours after plating, cells were transfected with PA-X variants tagged at the C terminus with eGFP or a 3xFlag tag following the protocols described above. 24 hours after transfection, cells were washed twice with phosphate buffer saline (PBS) and fixed with 4% paraformaldehyde. For PA-X-eGFP analysis, cells were then permeabilized in 0.1% Triton X100 and nuclei were stained with 1:10,000 dilution of Hoechst 3342 Fluorescent Stain (Fisher Scientific) for 10 minutes at room temperature. For PA-X-3xFlag staining, after fixation the cell membrane was stained by incubating cells for 10 minutes in 2.5 ug/ml wheat germ agglutinin (WGA) conjugated to Alexa Fluor™ 647 (Fisher Scientific, W32466) in PBS at room temperature in the dark. They were then permeabilized with 0.1% Triton X-100 in PBS for 10 minutes, blocked with 10% BSA in PBS for 30 minutes at 37°C, and incubated with 1:500 anti-Flag antibody (M2 clone, Sigma Aldrich F1804) in 3% BSA in PBS for 2 hrs at room temperature. Stained cells were washed in PBS and incubated with 5 ug/ml Goat anti-Mouse secondary antibody, Alexa Fluor™ 488 (ThermoFisher, A11029) in 3% BSA in PBS for 45 minutes at room temperature. Nuclei were stained with 1:10,000 dilution of Hoechst 3342 Fluorescent Stain (Fisher Scientific) for 10 minutes at room temperature. For all experiments, coverslips were then mounted on glass slides using ProLong Gold Antifade Mountant (Thermo Fisher). Images were taken with the Nikon A1R Confocal Microscope or the Leica Stellaris Confocal Microscope. To analyze PA-X subcellular localization, ImageJ was used to draw regions of interest for the nuclear and cytoplasmic compartments using Hoechst and autofluorescence or in the case of 3xFlag-tagged PA-X, WGA staining, respectively. The mean fluorescence intensity of eGFP/Flag signal in each compartment was then measured and the nuclear/cytoplasmic ratio of the mean intensity measurements was calculated. 10-15 cells per condition were analyzed for each replicate experiment.

### Protein collection and western blotting

Cell lysates were prepared using RIPA buffer (50 mM Tris-HCl pH 7.4, 1% NP-40 alternative, 0.5% sodium deoxycholate, 0.1% SDS, 150 mM NaCl, 2 mM EDTA) supplemented with 1X cOmplete EDTA-free protease inhibitor cocktail (Roche), 2 mM MgCl_2_, and 250 U/ml benzonase nuclease (Sigma-Aldrich). 100 μg of protein were loaded on Mini-PROTEAN® TGX™ Precast Protein Gels, 4–15% (10-well) or Mini-PROTEAN® TGX™ Precast Protein Gels, 4–20% (15 well) SDS-PAGE gels (Bio-Rad) and transferred onto polyvinylidene difluoride (PVDF) membranes (Millipore) using a Trans-Blot Turbo Transfer System (BioRad). Membranes were blocked with 5% milk in Tris buffer saline (TBS) with 0.1% Tween-20 (TBST). Western blots were performed at 4°C overnight using antibodies against GFP (ChromoTek pabg1-20, 1:500), myc tag (Cell Signaling Technologies #2276, 1:1000), Flag tag (Sigma-Aldrich F1804, 1:500), and β-tubulin (Cell Signaling Technologies #2128, 1:1000) diluted in 0.5% milk in TBST for anti-GFP, anti-Flag and anti-β-tubulin or 0.5% milk in PBS with 0.1% Tween-20 (PBST) for anti-myc. Secondary antibodies conjugated to horseradish peroxidase (HRP) (SouthernBiotech) were used at a 1:5000 dilution and a chemiluminescent signal was generated using the Pierce ECL Western Blotting Substrate or SuperSignal West Atto Ultimate Sensitivity Substrate (Thermo Fisher). Blots were then imaged using an iBright FL1000 Imaging System (iBright Firmware version 1.8.1).

### Mass spectrometry sample collection by immunoprecipitation

HEK293T cells were plated on 15 cm^2^ plates. 24 hours after plating, cells were transfected with 15 ug of pCR3.1 plasmids containing C-terminal eGFP-tagged catalytically inactive PA-X D108A or PA-X variants containing both the D108A mutation and the E2 mutations in 10% FBS in DMEM using jetPRIME transfection reagent (Polyplus). eGFP-tagged proteins were isolated from transfected cell lysates using GFP-Trap Magnetic Beads (ChromoTek) following the manufacturer’s protocol. Briefly, 24 hours post-transfection, cell lysates were prepared using RIPA buffer (10 mM Tris-HCl pH 7.5, 150 mM NaCl, 0.5 mM EDTA, 0.1% SDS, 1% Triton™ X-100, 1% deoxycholate) supplemented with 1X cOmplete EDTA-free protease inhibitor cocktail (Roche), 1 mM PMSF (G-Biosciences), 250 U/ml benzonase nuclease (Sigma-Aldrich), and 2.5 mM MgCl_2_ to ensure complete cell and nuclear lysis. Lysates were then diluted in Dilution buffer (10 mM Tris-HCl pH 7.5, 150 mM NaCl, 0.5 mM EDTA) supplemented with 1X cOmplete EDTA-free protease inhibitor cocktail (Roche) and 1 mM PMSF. Diluted lysates were added to GFP-Trap Magnetic beads equilibrated in Dilution buffer and rotated end-over-end at 4°C overnight. Beads were washed in Wash buffer (10 mM Tris-HCl pH 7.5, 150 mM NaCl, 0.05 % Nonidet™ P40 Substitute, 0.5 mM EDTA) and protein was eluted in 200 µl of Acidic elution buffer (200 mM glycine pH 2.5). The eluate was separated from the beads using a magnet and was neutralized by adding Neutralization buffer (1 M Tris pH 10.4) to 1/10th of the volume of the Acidic elution buffer.

### Mass spectrometry sample preparation using enzymatic “in liquid” digestion

Immunoprecitation eluates in Acidic elution buffer (200 µl) were diluted with 360 µl MilliQ-purified water, 140 µl trichloroacetic acid (TCA) and 700 µl acetone (10% TCA and 50% acetone vol:vol final concentration) and incubated on ice for 45 minutes to precipitate proteins. Precipitated proteins were collected by centrifugation for 10 minutes at room temperature at 16,000 xg and washed twice with cold acetone using the same centrifugation conditions. Protein pellets were air-dried briefly and re-solubilized in reducing buffer (20 μl of 8 M Urea in 50 mM NH_4_HCO_3_, pH 8.5) overnight at 4°C. To reduce disulfide bonds, 1.5 μl of 25 mM dithiothreitol (DTT) in 25 mM NH_4_HCO_3_ (pH 8.5) was added to samples, which were then incubated for 15 minutes at 56°C. Samples were cooled to room temperature. Cysteines were then alkylated by adding 1.8 μl of 55 mM chloroacetamide and incubating in the dark at room temperature for 15 minutes. The reaction was quenched with 4.8 μl of 25 mM DTT. Finally, 2.9 μl of lysyl endopeptidase solution (100 ng/μl LysC in 25 mM NH_4_HCO_3_, pH 8.5) (FujiFilm)) and 9 μl of 25 mM NH_4_HCO_3_ (pH 8.5) were added to a final volume of 40 µl and samples were incubated for 3 hours at 37°C for peptide digestion. The reaction was terminated by acidification with 2.5% trifluoroacetic acid (TFA) added to a 0.3% final concentration. Mass spectrometry sample preparation was carried out by the University of Wisconsin Biotechnology Center Mass Spectrometry Core Facility.

### Nano-liquid chromatography coupled with tandem mass spectrometry (NanoLC-MS/MS)

Digests were desalted using Pierce™ C18 SPE pipette tips (100 µl volume) per manufacturer protocol, eluted in 20 µl of 70/30/0.1% acetonitrile/H_2_O/TFA, dried to completion in a SpeedVac vacuum concentrator and finally reconstituted in 15 µl of 0.1% formic acid containing 2% acetonitrile. Peptides were analyzed by nanoLC-MS/MS using the Agilent 1100 Nanoflow system (Agilent) connected to hybrid linear ion trap-orbitrap mass spectrometer (LTQ-Orbitrap Elite™, Thermo Fisher Scientific) equipped with an EASY-Spray™ electrospray source (held at constant 45°C). Chromatography of peptides prior to mass spectral analysis was accomplished using capillary emitter column (PepMap® C18, 3 µM, 100 Å, 150×0.075 mm, Thermo Fisher Scientific) onto which 3 µl of extracted peptides were automatically loaded. NanoHPLC system delivered solvents A: 0.1% (v/v) formic acid, and B: 99.9% (v/v) acetonitrile, 0.1% (v/v) formic acid at 0.50 µL/min to load the peptides (over a 30 minute period) and 0.3 µl/min to elute peptides directly into the nano-electrospray with gradual gradient from 0% (v/v) B to 30% (v/v) B over 80 minutes and concluded with 5 minute fast gradient from 30% (v/v) B to 50% (v/v) B at which time a 5 minute flash-out from 50-95% (v/v) B took place. As peptides eluted from the HPLC-column/electrospray source survey MS scans were acquired in the Orbitrap with a resolution of 120,000 followed by CID-type MS/MS fragmentation of 30 most intense peptides detected in the MS1 scan from 350 to 1800 m/z; redundancy was limited by dynamic exclusion. Mass spectrometry analysis was carried out by the University of Wisconsin Biotechnology Center Mass Spectrometry Core Facility.

### Mass spectrometry data analysis

LTQ-Orbitrap Elite-acquired raw MS/MS data files were converted to mgf file format using MSConvert (ProteoWizard: Open Source Software for Rapid Proteomics Tools Development). Resulting mgf files were used to search against user defined human database (Homo sapiens UP000005640 UniProt reference proteome, 03/08/2024 download containing 82,602 protein entries) plus nine PA-X mutant sequences along with a common Repository of Adventitious Proteins (cRAP), a database of common lab contaminants (116 total entries), using in-house Mascot search engine 3.0.0 (Matrix Science). Carbamidomethylation of cysteines was set as a fixed modification, while N-terminal acetylation, methionine oxidation, and deamidation of asparagine and glutamine were set as variable modifications. Peptide mass tolerance was set at 10 parts per million and fragment mass at 0.6 Da. Peptide and protein annotations, significance of identification, and spectral based quantification was done with Scaffold software (version 5.0.1, Proteome Software Inc., Portland, OR). Peptide identifications were accepted if they could be established at greater than 90.0% probability to achieve a false discovery rate (FDR) less than 1.0% by the PeptideProphet algorithm(74) with Scaffold delta-mass correction. The ProteinProphet algorithm(75) grouped peptides by their corresponding protein(s). Protein identifications were accepted if they could be established at greater than 99.0% probability by the ProteinProphet algorithm to achieve an FDR less than 1.0% and contained at least 2 identified peptides. Proteins that contained similar peptides and could not be differentiated based on MS/MS analysis alone were grouped to satisfy the principles of parsimony. Mass spectrometry data analysis was carried out by the University of Wisconsin Biotechnology Center Mass Spectrometry Core Facility.

### 5’ rapid amplification of cDNA ends (RACE)

RNA was extracted from cells using the Quick-RNA miniprep kit (Zymo Research) following the manufacturer’s protocol and treated with Turbo DNase. After phenol/chloroform extraction from the DNase reaction, the RACE adapter (sequences below) was ligated to 6 μg RNA using T4 RNA ligase (Ambion) at 25°C for 2 hrs. MMLV RT (Invitrogen) was then used to reverse transcribe ligated RNA to cDNA following the manufacturer’s protocol. Fragments of interest were amplified by Taq DNA polymerase (New England Biolabs) using forward primers annealing to the RACE adapter and reverse primers annealing to the transfected reporter construct (sequences below(6, 68)). PCR products were separated on a 2% agarose gel containing HydraGreen safe DNA dye (ACTGene) and visualized using the iBright FL1000 imager system. DNA was extracted from gel bands at expected sizes and Sanger sequenced to confirm their identities.

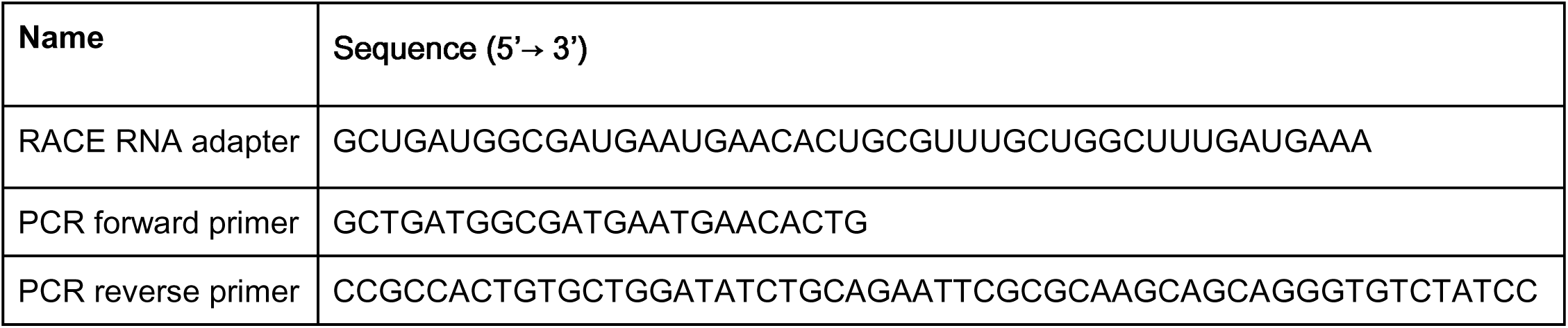

### Quantification and statistical analysis

Data plotted on graphs represent three or more independent biological replicates, as shown by the individual data points included in all graphs. Images shown are also representative of three or more independent biological replicates. Mass spectrometry analyses were carried out on samples collected in two separate experiments. Peptide counts were aggregated and plotted using stacked bar graphs as a percentage of total peptides detected. For box and whisker plots, the boxes extend from the 25th to 75th percentile of values, horizontal lines denote median values, and whiskers are plotted from the minimum to maximum values. Floating bar plots were used instead of box and whisker plots when n=3 (Fig. 3E, 4B), with boxes that extend from the minimum to maximum values and horizontal lines denoting median values. For the scatter plots (Fig. 1E, 2D, 3D, 4D), horizontal lines denote the mean and error bars denote the standard deviation. For multiple comparisons, one-way analysis of variance (ANOVA) followed by Dunnett’s multiple comparison test or two-way ANOVA followed by uncorrected Fisher’s LSD multiple comparison test were used, as indicated in the figure legends. Overall significance was defined as < 0.05. Where indicated, levels of significance are denoted as follows: ns = p > 0.05, * = p < 0.05, ** = p < 0.01, *** = p < 0.001, **** = p < 0.0001. Plotting and statistical analyses were performed using GraphPad Prism (v10.5.0). Several experiments were conducted in parallel but are plotted separately in the manuscript. Therefore, in Figures 1, 2, and 3, control conditions (WT PA-X, eGFP, or PA-X D108A) include the same data and are identical.

## Acknowledgements

We thank Drs C. McCormick, D. Khaperskyy, B. Glaunsinger, G. Dreyfuss, and R. Webby for constructs. We thank Drs C. McCormick, D. Khaperskyy, A. Mehle, and members of the Gaglia laboratory for feedback and advice. We thank the Tufts Center for Neuroscience Research Imaging and Cell Analysis Core and Dr. K. Majumder for technical support and assistance on the confocal microscopes used in this study. We thank the University of Wisconsin Biotechnology Center Mass Spectrometry Core Facility for technical guidance and performing the LC/MS/MS used in this study.

## Figure legends

## References

1. Jagger BW, Wise HM, Kash JC, Walters K-A, Wills NM, Xiao Y-L, Dunfee RL, Schwartzman LM, Ozinsky A, Bell GL, Dalton RM, Lo A, Efstathiou S, Atkins JF, Firth AE, Taubenberger JK, Digard P. 2012. An Overlapping Protein-Coding Region in Influenza A Virus Segment 3 Modulates the Host Response. Science 337:199–204.

2. Hayashi T, MacDonald LA, Takimoto T. 2015. Influenza A Virus Protein PA-X Contributes to Viral Growth and Suppression of the Host Antiviral and Immune Responses. J Virol 89:6442–6452.

3. Chaimayo C, Dunagan M, Hayashi T, Santoso N, Takimoto T. 2018. Specificity and functional interplay between influenza virus PA-X and NS1 shutoff activity. PLOS Pathogens 14:e1007465.

4. Khaperskyy DA, Schmaling S, Larkins-Ford J, McCormick C, Gaglia MM. 2016. Selective Degradation of Host RNA Polymerase II Transcripts by Influenza A Virus PA-X Host Shutoff Protein. PLoS Pathog 12:e1005427.

5. Gaucherand L, Porter BK, Levene RE, Price EL, Schmaling SK, Rycroft CH, Kevorkian Y, McCormick C, Khaperskyy DA, Gaglia MM. 2019. The Influenza A Virus Endoribonuclease PA-X Usurps Host mRNA Processing Machinery to Limit Host Gene Expression. Cell Reports 27:776–792.e7.

6. Gaucherand L, Iyer A, Gilabert I, Rycroft CH, Gaglia MM. 2023. Cut site preference allows influenza A virus PA-X to discriminate between host and viral mRNAs. 7. Nat Microbiol 8:1304–1317.

7. Hayashi T, Chaimayo C, McGuinness J, Takimoto T. 2016. Critical Role of the PA-X C-Terminal Domain of Influenza A Virus in Its Subcellular Localization and Shutoff Activity. Journal of Virology 90:7131–7141.

8. Levene RE, Shrestha SD, Gaglia MM. 2021. The influenza A virus host shutoff factor PA-X is rapidly turned over in a strain-specific manner. J Virol 10.1128/JVI.02312-20.

9. Oishi K, Yamayoshi S, Kozuka-Hata H, Oyama M, Kawaoka Y. 2018. N-Terminal Acetylation by NatB Is Required for the Shutoff Activity of Influenza A Virus PA-X. Cell Reports 24:851–860.

10. Brown JL, Roberts WK. 1976. Evidence that approximately eighty per cent of the soluble proteins from Ehrlich ascites cells are Nalpha-acetylated. Journal of Biological Chemistry 251:1009–1014.

11. Arnesen T, Van Damme P, Polevoda B, Helsens K, Evjenth R, Colaert N, Varhaug JE, Vandekerckhove J, Lillehaug JR, Sherman F, Gevaert K. 2009. Proteomics analyses reveal the evolutionary conservation and divergence of N-terminal acetyltransferases from yeast and humans. Proc Natl Acad Sci U S A 106:8157–8162.

12. Aksnes H, Hole K, Arnesen T. 2015. Molecular, cellular, and physiological significance of N-terminal acetylation. Int Rev Cell Mol Biol 316:267–305.

13. Aksnes H, Drazic A, Marie M, Arnesen T. 2016. First Things First: Vital Protein Marks by N-Terminal Acetyltransferases. Trends in Biochemical Sciences 41:746–760.

14. Starheim KK, Gevaert K, Arnesen T. 2012. Protein N-terminal acetyltransferases: when the start matters. Trends in Biochemical Sciences 37:152–161.

15. Ree R, Varland S, Arnesen T. 2018. Spotlight on protein N-terminal acetylation. Experimental & Molecular Medicine 50:90.

16. Arnesen T. 2011. Towards a Functional Understanding of Protein N-Terminal Acetylation. PLOS Biology 9:e1001074.

17. Myklebust LM, Van Damme P, Støve SI, Dörfel MJ, Abboud A, Kalvik TV, Grauffel C, Jonckheere V, Wu Y, Swensen J, Kaasa H, Liszczak G, Marmorstein R, Reuter N, Lyon GJ, Gevaert K, Arnesen T. 2015. Biochemical and cellular analysis of Ogden syndrome reveals downstream Nt-acetylation defects. Hum Mol Genet 24:1956–1976.

18. Hwang C-S, Shemorry A, Varshavsky A. 2010. N-Terminal Acetylation of Cellular Proteins Creates Specific Degradation Signals. Science 327:973–977.

19. Shemorry A, Hwang C-S, Varshavsky A. 2013. Control of protein quality and stoichiometries by N-terminal acetylation and the N-end rule pathway. Mol Cell 50:540– 551.

20. Xu F, Huang Y, Li L, Gannon P, Linster E, Huber M, Kapos P, Bienvenut W, Polevoda B, Meinnel T, Hell R, Giglione C, Zhang Y, Wirtz M, Chen S, Li X. 2015. Two N-Terminal Acetyltransferases Antagonistically Regulate the Stability of a Nod-Like Receptor in Arabidopsis. Plant Cell 27:1547–1562.

21. Sheikh TI, de Paz AM, Akhtar S, Ausió J, Vincent JB. 2017. MeCP2_E1 N-terminal modifications affect its degradation rate and are disrupted by the Ala2Val Rett mutation. Hum Mol Genet 26:4132–4141.

22. Holmes WM, Mannakee BK, Gutenkunst RN, Serio TR. 2014. Loss of amino-terminal acetylation suppresses a prion phenotype by modulating global protein folding. 1. Nature Communications 5:4383.

23. Trexler AJ, Rhoades E. 2012. N-terminal acetylation is critical for forming α-helical oligomer of α-synuclein. Protein Sci 21:601–605.

24. Bartels T, Choi JG, Selkoe DJ. 2011. α-Synuclein occurs physiologically as a helically folded tetramer that resists aggregation. Nature 477:107–110.

25. Scott DC, Monda JK, Bennett EJ, Harper JW, Schulman BA. 2011. N-Terminal Acetylation Acts as an Avidity Enhancer Within an Interconnected Multiprotein Complex. Science 334:674–678.

26. Monda JK, Scott DC, Miller DJ, Lydeard J, King D, Harper JW, Bennett EJ, Schulman BA. 2013. Structural Conservation of Distinctive N-terminal Acetylation-Dependent Interactions across a Family of Mammalian NEDD8 Ligation Enzymes. Structure 21:42–53.

27. Yang D, Fang Q, Wang M, Ren R, Wang H, He M, Sun Y, Yang N, Xu R-M. 2013. Nα-acetylated Sir3 stabilizes the conformation of a nucleosome-binding loop in the BAH domain. 9. Nature Structural & Molecular Biology 20:1116–1118.

28. Arnaudo N, Fernández IS, McLaughlin SH, Peak-Chew SY, Rhodes D, Martino F. 2013. The N-terminal acetylation of Sir3 stabilizes its binding to the nucleosome core particle. 9. Nature Structural & Molecular Biology 20:1119–1121.

29. Singer JM, Shaw JM. 2003. Mdm20 protein functions with Nat3 protein to acetylate Tpm1 protein and regulate tropomyosin–actin interactions in budding yeast. Proc Natl Acad Sci U S A 100:7644–7649.

30. Coulton AT, East DA, Galinska-Rakoczy A, Lehman W, Mulvihill DP. 2010. The recruitment of acetylated and unacetylated tropomyosin to distinct actin polymers permits the discrete regulation of specific myosins in fission yeast. J Cell Sci 123:3235–3243.

31. Forte GMA, Pool MR, Stirling CJ. 2011. N-Terminal Acetylation Inhibits Protein Targeting to the Endoplasmic Reticulum. PLOS Biology 9:e1001073.

32. Setty SRG, Strochlic TI, Tong AHY, Boone C, Burd CG. 2004. Golgi targeting of ARF-like GTPase Arl3p requires its Nα-acetylation and the integral membrane protein Sys1p. 5. Nat Cell Biol 6:414–419.

33. Behnia R, Panic B, Whyte JRC, Munro S. 2004. Targeting of the Arf-like GTPase Arl3p to the Golgi requires N-terminal acetylation and the membrane protein Sys1p. 5. Nat Cell Biol 6:405–413.

34. Behnia R, Barr FA, Flanagan JJ, Barlowe C, Munro S. 2007. The yeast orthologue of GRASP65 forms a complex with a coiled-coil protein that contributes to ER to Golgi traffic. J Cell Biol 176:255–261.

35. Shi M, Jagger BW, Wise HM, Digard P, Holmes EC, Taubenberger JK. 2012. Evolutionary Conservation of the PA-X Open Reading Frame in Segment 3 of Influenza A Virus. Journal of Virology 86:12411–12413.

36. Oishi K, Yamayoshi S, Kawaoka Y. 2015. Mapping of a Region of the PA-X Protein of Influenza A Virus That Is Important for Its Shutoff Activity. J Virol 89:8661–8665.

37. Goetze S, Qeli E, Mosimann C, Staes A, Gerrits B, Roschitzki B, Mohanty S, Niederer EM, Laczko E, Timmerman E, Lange V, Hafen E, Aebersold R, Vandekerckhove J, Basler K, Ahrens CH, Gevaert K, Brunner E. 2009. Identification and Functional Characterization of N-Terminally Acetylated Proteins in Drosophila melanogaster. PLOS Biology 7:e1000236.

38. Kelley JB, Paschal BM. 2019. Fluorescence-based Quantification of Nucleocytoplasmic Transport. Methods 157:106–114.

39. Schindelin J, Arganda-Carreras I, Frise E, Kaynig V, Longair M, Pietzsch T, Preibisch S, Rueden C, Saalfeld S, Schmid B, Tinevez J-Y, White DJ, Hartenstein V, Eliceiri K, Tomancak P, Cardona A. 2012. Fiji: an open-source platform for biological-image analysis. 7. Nature Methods 9:676–682.

40. Adam SA, Lobl TJ, Mitchell MA, Gerace L. 1989. Identification of specific binding proteins for a nuclear location sequence. 6204. Nature 337:276–279.

41. Aksnes H, Van Damme P, Goris M, Starheim KK, Marie M, Støve SI, Hoel C, Kalvik TV, Hole K, Glomnes N, Furnes C, Ljostveit S, Ziegler M, Niere M, Gevaert K, Arnesen T. 2015. An Organellar Nα-Acetyltransferase, Naa60, Acetylates Cytosolic N Termini of Transmembrane Proteins and Maintains Golgi Integrity. Cell Reports 10:1362–1374.

42. Wingfield P. 2017. N-Terminal Methionine Processing. Curr Protoc Protein Sci 88:6.14.1–6.14.3.

43. Van Damme P. 2021. Charting the N-Terminal Acetylome: A Comprehensive Map of Human NatA Substrates. Int J Mol Sci 22:10692.

44. CDC. 2024. Types of Influenza Viruses. Influenza (Flu). https://www.cdc.gov/flu/about/viruses-types.html. Retrieved 14 July 2025.

45. Thompson WW, Weintraub E, Dhankhar P, Cheng P, Brammer L, Meltzer MI, Bresee JS, Shay DK. 2009. Estimates of US influenza-associated deaths made using four different methods. Influenza Other Respir Viruses 3:37–49.

46. Lanz C, Schotsaert M, Magnus C, Karakus U, Hunziker A, Sempere Borau M, Martínez-Romero C, Spieler EE, Günther SC, Moritz E, Hale BG, Trkola A, García-Sastre A, Stertz S. 2021. IFITM3 incorporation sensitizes influenza A virus to antibody-mediated neutralization. J Exp Med 218:e20200303.

47. Xia C, Vijayan M, Pritzl CJ, Fuchs SY, McDermott AB, Hahm B. 2016. Hemagglutinin of Influenza A Virus Antagonizes Type I Interferon (IFN) Responses by Inducing Degradation of Type I IFN Receptor 1. J Virol 90:2403–2417.

48. Zhang B, Xu S, Liu M, Wei Y, Wang Q, Shen W, Lei C-Q, Zhu Q. The nucleoprotein of influenza A virus inhibits the innate immune response by inducing mitophagy. Autophagy 19:1916–1933.

49. Wang S, Li Z, Chen Y, Gao S, Qiao J, Liu H, Song H, Ao D, Sun X. 2023. ARIH1 inhibits influenza A virus replication and facilitates RIG-I dependent immune signaling by interacting with SQSTM1/p62. Virol J 20:58.

50. Jørgensen SE, Christiansen M, Ryø LB, Gad HH, Gjedsted J, Staeheli P, Mikkelsen JG, Storgaard M, Hartmann R, Mogensen TH. 2018. Defective RNA sensing by RIG-I in severe influenza virus infection. Clin Exp Immunol 192:366–376.

51. Petrich A, Chiantia S. 2023. Influenza A Virus Infection Alters Lipid Packing and Surface Electrostatic Potential of the Host Plasma Membrane. 9. Viruses 15:1830.

52. Rodriguez A, Pérez-González A, Nieto A. 2007. Influenza virus infection causes specific degradation of the largest subunit of cellular RNA polymerase II. J Virol 81:5315–5324.

53. Liu X, Xu F, Ren L, Zhao F, Huang Y, Wei L, Wang Y, Wang C, Fan Z, Mei S, Song J, Zhao Z, Cen S, Liang C, Wang J, Guo F. 2021. MARCH8 inhibits influenza A virus infection by targeting viral M2 protein for ubiquitination-dependent degradation in lysosomes. Nat Commun 12:4427.

54. Yu L, Jiang Y, Rang H, Wang X, Cai Y, Yan H, Wu S, Lan K. 2025. Restriction of influenza A virus replication by host DCAF7-CRL4B axis. Journal of Virology 99:e00133–25.

55. Bougon J, Kadijk E, Gallot-Lavallee L, Curtis BA, Landers M, Archibald JM, Khaperskyy DA. Influenza A virus NS1 effector domain is required for PA-X-mediated host shutoff in infected cells. J Virol 98:e01901–23.

56. Khaperskyy DA, Emara MM, Johnston BP, Anderson P, Hatchette TF, McCormick C. 2014. Influenza A Virus Host Shutoff Disables Antiviral Stress-Induced Translation Arrest. PLoS Pathog 10:e1004217.

57. Poon IKH, Jans DA. 2005. Regulation of nuclear transport: central role in development and transformation? Traffic 6:173–186.

58. Bartko JC, Li Y, Sun G, Halterman MW. 2020. Phosphorylation within the bipartite NLS alters the localization and toxicity of the ER stress response factor DDIT3/CHOP. Cell Signal 74:109713.

59. Hofweber M, Hutten S, Bourgeois B, Spreitzer E, Niedner-Boblenz A, Schifferer M, Ruepp M-D, Simons M, Niessing D, Madl T, Dormann D. 2018. Phase Separation of FUS Is Suppressed by Its Nuclear Import Receptor and Arginine Methylation. Cell 173:706–719.e13.

60. Zakaryan RP, Gehring H. 2006. Identification and characterization of the nuclear localization/retention signal in the EWS proto-oncoprotein. J Mol Biol 363:27–38.

61. Mallet P-L, Bachand F. 2013. A proline-tyrosine nuclear localization signal (PY-NLS) is required for the nuclear import of fission yeast PAB2, but not of human PABPN1. Traffic 14:282–294.

62. Bannister AJ, Miska EA, Görlich D, Kouzarides T. 2000. Acetylation of importin-alpha nuclear import factors by CBP/p300. Curr Biol 10:467–470.

63. Dikiy I, Eliezer D. 2014. N-terminal acetylation stabilizes N-terminal helicity in lipid– and micelle-bound α-synuclein and increases its affinity for physiological membranes. J Biol Chem 289:3652–3665.

64. Coombs KM, Berard A, Xu W, Krokhin O, Meng X, Cortens JP, Kobasa D, Wilkins J, Brown EG. 2010. Quantitative Proteomic Analyses of Influenza Virus-Infected Cultured Human Lung Cells. Journal of Virology 84:10888–10906.

65. Haas KM, McGregor MJ, Bouhaddou M, Polacco BJ, Kim E-Y, Nguyen TT, Newton BW, Urbanowski M, Kim H, Williams MAP, Rezelj VV, Hardy A, Fossati A, Stevenson EJ, Sukerman E, Kim T, Penugonda S, Moreno E, Braberg H, Zhou Y, Metreveli G, Harjai B, Tummino TA, Melnyk JE, Soucheray M, Batra J, Pache L, Martin-Sancho L, Carlson-Stevermer J, Jureka AS, Basler CF, Shokat KM, Shoichet BK, Shriver LP, Johnson JR, Shaw ML, Chanda SK, Roden DM, Carter TC, Kottyan LC, Chisholm RL, Pacheco JA, Smith ME, Schrodi SJ, Albrecht RA, Vignuzzi M, Zuliani-Alvarez L, Swaney DL, Eckhardt M, Wolinsky SM, White KM, Hultquist JF, Kaake RM, García-Sastre A, Krogan NJ. 2023. Proteomic and genetic analyses of influenza A viruses identify pan-viral host targets. 1. Nat Commun 14:6030.

66. Younis I, Berg M, Kaida D, Dittmar K, Wang C, Dreyfuss G. 2010. Rapid-Response Splicing Reporter Screens Identify Differential Regulators of Constitutive and Alternative Splicing. Mol Cell Biol 30:1718–1728.

67. Hoffmann E, Krauss S, Perez D, Webby R, Webster RG. 2002. Eight-plasmid system for rapid generation of influenza virus vaccines. Vaccine 20:3165–3170.

68. Gaglia MM, Rycroft CH, Glaunsinger BA. 2015. Transcriptome-Wide Cleavage Site Mapping on Cellular mRNAs Reveals Features Underlying Sequence-Specific Cleavage by the Viral Ribonuclease SOX. PLoS Pathog 11:e1005305.

69. Matrosovich M, Matrosovich T, Garten W, Klenk H-D. 2006. New low-viscosity overlay medium for viral plaque assays. Virol J 3:63.

70. Kuchipudi SV, Tellabati M, Nelli RK, White GA, Perez BB, Sebastian S, Slomka MJ, Brookes SM, Brown IH, Dunham SP, Chang K-C. 2012. 18S rRNA is a reliable normalisation gene for real time PCR based on influenza virus infected cells. Virol J 9:230.

71. Abernathy E, Gilbertson S, Alla R, Glaunsinger B. 2015. Viral Nucleases Induce an mRNA Degradation-Transcription Feedback Loop in Mammalian Cells. Cell Host & Microbe 18:243–253.

72. Hu T, Zhang C, Tang Q, Su Y, Li B, Chen L, Zhang Z, Cai T, Zhu Y. 2013. Variant G6PD levels promote tumor cell proliferation or apoptosis via the STAT3/5 pathway in the human melanoma xenograft mouse model. BMC Cancer 13:251.

73. Slaine PD, Kleer M, Duguay BA, Pringle ES, Kadijk E, Ying S, Balgi A, Roberge M, McCormick C, Khaperskyy DA. 2021. Thiopurines Activate an Antiviral Unfolded Protein Response That Blocks Influenza A Virus Glycoprotein Accumulation. J Virol 95:e00453–21.

74. Keller A, Nesvizhskii AI, Kolker E, Aebersold R. 2002. Empirical Statistical Model To Estimate the Accuracy of Peptide Identifications Made by MS/MS and Database Search. Anal Chem 74:5383–5392.

75. Nesvizhskii AI, Keller A, Kolker E, Aebersold R. 2003. A Statistical Model for Identifying Proteins by Tandem Mass Spectrometry. Anal Chem 75:4646–4658.

